# Simulating the emergence of the Visual Word Form Area: Recycling a convolutional neural network for reading

**DOI:** 10.1101/2021.02.15.431235

**Authors:** T. Hannagan, A. Agrawal, L. Cohen, S. Dehaene

## Abstract

The visual word form area (VWFA) is a region of human inferotemporal cortex that emerges at a fixed location in occipitotemporal cortex during reading acquisition, and systematically responds to written words in literate individuals. According to the neuronal recycling hypothesis, this region arises through the repurposing, for letter recognition, of a subpart of the ventral visual pathway initially involved in face and object recognition. Furthermore, according to the biased connectivity hypothesis, its universal localization is due to pre-existing connections from this subregion to areas involved in spoken language processing. Here, we evaluate those hypotheses in an explicit computational model. We trained a deep convolutional neural network of the ventral visual pathway, first to categorize pictures, and then to recognize written words invariantly for case, font and size. We show that the model can account for many properties of the VWFA, particularly when a subset of units possesses a biased connectivity to word output units. The network develops a sparse, invariant representation of written words, based on a restricted set of reading-selective units. Their activation mimics several properties of the VWFA, and their lesioning causes a reading-specific deficit. Our simulation fleshes out the neuronal recycling hypothesis, and make several testable predictions concerning the neural code for written words.

## Introduction

Reading acquisition relies on the development of a novel interface between vision and language, in charge of efficiently identifying letters and their ordering [1,2]. This orthographic analysis then feeds the language systems supporting semantics and phonology. Over the past 20 years, some basic features of this interface have been put to light. A specific region of the left ventral occipitotemporal cortex (VOT), which was labelled the Visual Word Form Area (VWFA), is present at a similar location in the brain of every literate subject and is thought to underlie orthographic coding [3]. Functional brain imaging has uncovered a host of functional features of the VWFA, for example tuning to familiar vs. unknown alphabets [4,5], partial invariance for retinal location [3,6], invariance for upper/lower case [7,8], or sensitivity to the frequency of word occurrence [9,10]. Nevertheless, how this region becomes specialized for written words, or even whether it does so, remain highly controversial issues [11–14].

Current evidence suggests that the VWFA site owes its functional specialization to a combination of two factors. First, according to the neuronal recycling hypothesis, reading pre-empts and repurposes part of the large region of ventral visual cortex that participates in visual object recognition [11,15–17]. The specific region involved may not only possess a generic architecture for invariant visual recognition, but also a bottom-up sensitivity to some of the shape features relevant to word recognition, such as a preference for high-resolution foveal inputs [18,19], line junctions [20] and mid-level geometrical features [21]. Because these properties are widespread in both hemispheres, however, a second hypothesis may be needed to explain the narrow, reproducible location of the VWFA in the depth of the left infero-temporal sulcus. According to the biased connectivity hypothesis, this left-hemispheric site exhibits a pre-existing biased connectivity, or “connectivity fingerprints” with distant language areas [22–27]. Indeed, in agreement with this idea, the precise location of the VWFA in 8-years-old readers can be predicted from their long-distance anatomical connectivity to other brain areas at five years of age, before they learned to read [27].

In the present work, we assess to what extent a minimal computational model of those two hypotheses may suffice to account for the emergence of the VWFA during reading acquisition. Specifically, we simulate a deep neural network whose architecture was not designed for reading, but is inspired from that of the ventral visual cortex and which was shown to provide a good fit to both behavioral and electrophysiological observations on face and object recognition [28]. We examinee what happens when this network, after being trained to identify pictures of generic object categories, is further taught to identify written words, with and without biased connections to output lexical units.

### Aims of the present study

Our work had two aims. First, we wanted to see if we could reproduce a list of experimentally observed properties of the VWFA, namely

- Emergence, after training, of a localized patch of neurons specialized for words as opposed to other stimuli such as faces or objects [11]
- Recycling of units with modest prior involvement for objects and faces prior to reading acquisition [11]
- Invariance for word size, case and font [3,6–8]
- Monotonically increasing response to a hierarchy of letter strings that increasingly approximate the statistics of words in the learned script [29–31]
- Sudden loss of reading abilities (‘pure alexia’) when this patch of cortex is lesioned [32– 34], with preserved recognition of other visual categories.

A failure to capture some of these properties with a simple feedforward convolutional neural network would be interesting inasmuch as it may point to the need for additional properties, for instance recurrent and/or top-down connections [12,14,31,35].

Our second goal was to see if we could predict, in anticipation of future experiments, some of the properties of the neural code for written words. It is currently controversial whether neurons in the reading pathway are specialized for whole words [36], frequent pairs of letters (‘bigrams’) [37,38], graphemes that map onto phonemes [39], or individual letters at a specific location [40,41]. Indeed, these possibilities are not mutually exclusive, and multiple codes may coexist, perhaps in different pathways, to support different tasks such as comprehension versus reading aloud [39,42]. While it is currently nearly impossible to visualize single neurons in the reading pathway in humans, this has been achieved in a non-human primate monkey model [43], and advances in intracranial recordings may soon make it possible in the human brain [31,44]. Using as few prior hypotheses as possible, we describe how an artificial neural network encodes visual words, in the hope that its predicted tuning curves may soon become empirically testable.

## The model

### Architecture

Our model is based on CORnet-Z, the simplest network in the CORnet family [28]. These convolutional neural networks (CNN) all share a common design of 4 spatially-organized modules meant to represent V1, V2, V4 and IT areas in the ventral visual pathway (topological modules hereafter), capped by a non-topological decoding structure (dense layer in Fig. 1). We built three variants of CORnet-Z networks. The illiterate network was trained only on the visual images in ImageNet (ILSVRC-2012 dataset, about 1 300 000 image exemplars distributed over 1000 classes such as dogs, cars etc). Two literate networks were additionally trained to recognize words. To this end, we generated 1 300 000 word exemplars distributed over 1000 “word classes”, where each word class actually corresponded to a single word, while exemplars varied in location, size, font and scale. The literate networks thus had 1000 additional output units (see Fig. 1). In the unbiased literate network, their afferent connections were distributed across all units in the dense layer. In the biased literate network, to simulate the hypothesis of a biased connectivity from a subregion of visual cortex to language areas, we restricted the output units’ afferent connections to a subset of 49 units in the dense layer. These biased units will be dubbed “Dense+” hereafter (violet units in Figure 1C), in opposition to unbiased units (“Dense-”, white units in Figure 1A, 1B, 1C). The code for training the networks is available online at https://github.com/THANNAGA/Origins-of-VWFA along with pre-trained models for all conditions.

**Fig 1.**
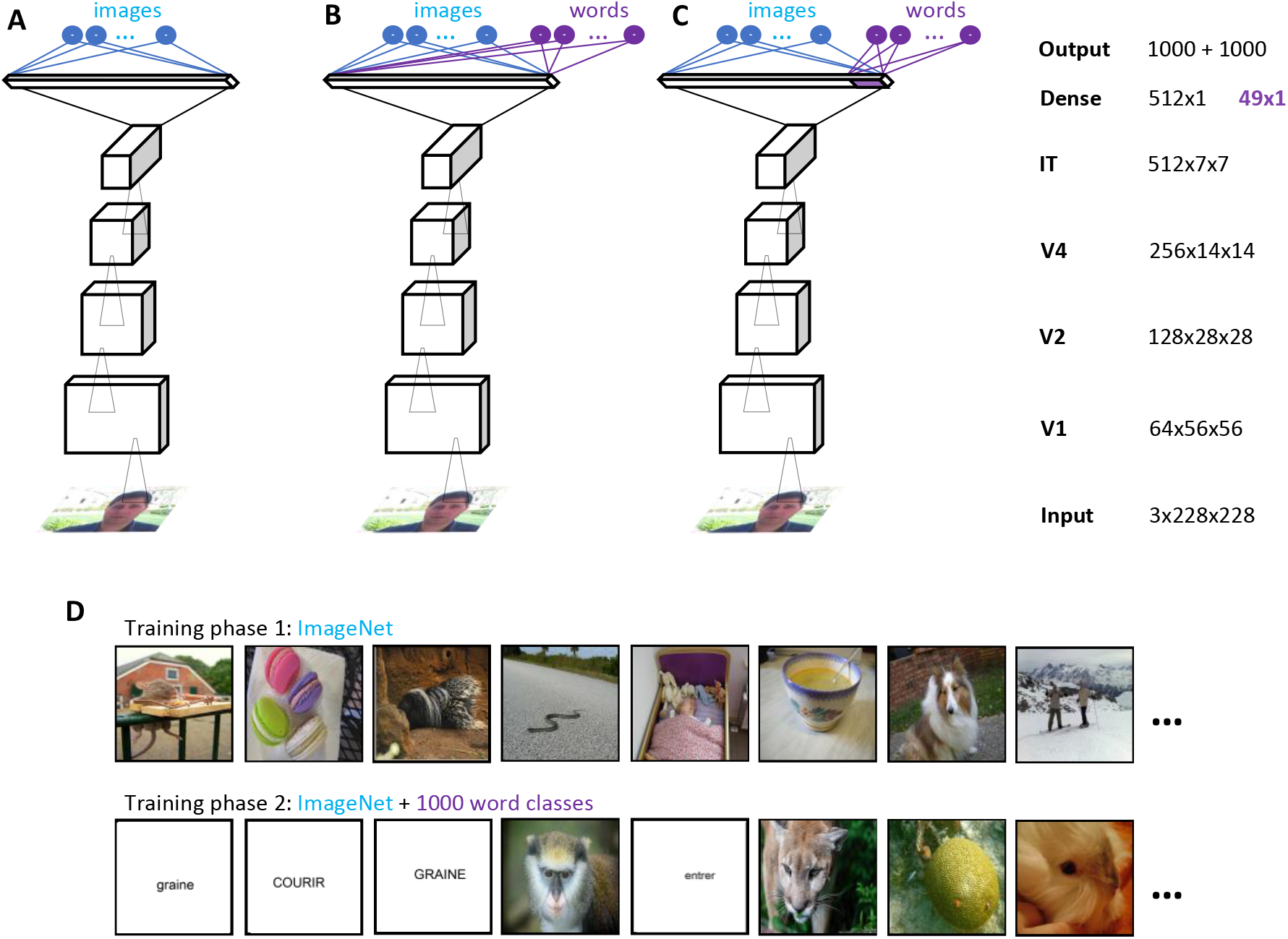
Network architectures and training schemes. Testing the biased-connectivity hypothesis using the CORnet convolution neural network model of the ventral visual pathway. A, Illiterate model: network trained only on ImageNet. B, Unbiased literate model: network first trained on ImageNet, then on ImageNet and words, with a new set of fully connected output word units. C, Biased literate model: network trained on ImageNet, then on ImageNet and words, with biased connectivity between a subset of units in the dense layer, and the output word units. A literate model before the introduction of words is referred to as “pre-literate”. D, Examples of ImageNet and word stimuli illustrating the training sets in phases 1 and 2.

### Stimuli

We used the training and validation sets from ImageNet, amounting to 1000 image classes, each with about 1300 training exemplars and 50 validation exemplars. Images were rescaled to 228×228 pixels before presentation. In addition to ImageNet, we also generated written word stimuli. All word exemplars were black-on-white images of dimension 228×228 pixels. The word set was selected from 1750 high-frequency French words known by 4 years old, as listed by the French Academy of Amiens (http://www.langage-en-maternelle.fr/PDF/1750_mots_quatre_ans.pdf). Among this list, we randomly selected 1000 words whose length ranged between 3 and 8 letters, without accents. Each word was considered as a class on its own, and we generated 1300 training exemplars of each word, varying in size, font, location and case, and 50 validation exemplars. In the training set, character size (scale) varied randomly between 40 and 80 pixels, fonts were either ‘Arial’ or ‘Times new roman’, and case was either upper or lower. Words were randomly shifted away from a central presentation, with shifts ranging uniformly from −50 to +50 pixels along the x-axis, and −30 to +30 pixels along the y-axis. Exemplars were generated with a uniform probability for all of these variables. In the validation set, the 50 exemplars for each word were generated in the same way, except that the fonts were randomly chosen between ‘Calibri’, ‘Courier’ and ‘Comic sans’.

### Training

Deep learning models typically involve a single training phase on a single large dataset, whereas children learn to identify faces and objects long before they acquire reading. To simulate reading acquisition as the partial recycling of prior visual recognition abilities, we trained the networks in two stages: an “image” phase (phase 1) followed by an “image + word” phase (phase 2). Word units were only added to the output layer of the network during the latter phase. Throughout phases 1 and 2, the networks were trained with Stochastic Gradient Descent on a categorical cross-entropy loss, using a linear learning rate scheduling (see Supplementary Material).

Neural networks are notoriously prone to the phenomenon of catastrophic interference, whereby learning of a new task, or even on a new dataset, can spectacularly deteriorate previously stored knowledge. This well-studied effect can be alleviated in a number of ways, included interleaved learning and the use of dropout [45,46]. We used interleaved learning: during phase 2, output word units were added to the network and the new training set was obtained by concatenating and shuffling the original generic image set and the new word set. The sets were matched for number of classes and exemplars, and the presentation probabilities were 50% images and 50% words. An illiterate control network was trained only on ImageNet, for as many epochs as the literate networks.

While it is common to freeze a network’s lower-level weights during transfer of learning to new datasets or tasks, here we trained the full network during the first and second stages, mindful of the fact that an effect of literacy has been reported as early as V1 in the visual pathway [17]. Finally, deep learning models trained on classification usually also augment the training set by performing various visual transforms which do no impact on the semantics of the image, such as cropping, resizing and mirroring. We included such transformations in our training set, but because words are not normally seen in mirror form, and for the sake of a fair comparison of network performance between words and pictures, we excluded mirror transforms for all classes during training.

## Results

### Training and recognition performance

For each network and at each epoch of training, we computed the top-1 accuracy on the validation set, i.e. the proportion of correctly classified exemplars in the validation set. Figure 2A shows that performance for all networks on ImageNet (light blue curves) was on par with earlier reports for CorNet-Z. However, despite the significant variability in script, case, font and size, words were learned better and quicker than images. In both literate networks and within 5 epochs, word accuracies on the validation set rose to 80% (dark blue curves), i.e. more than twice as high as for images.

**Fig 2.**
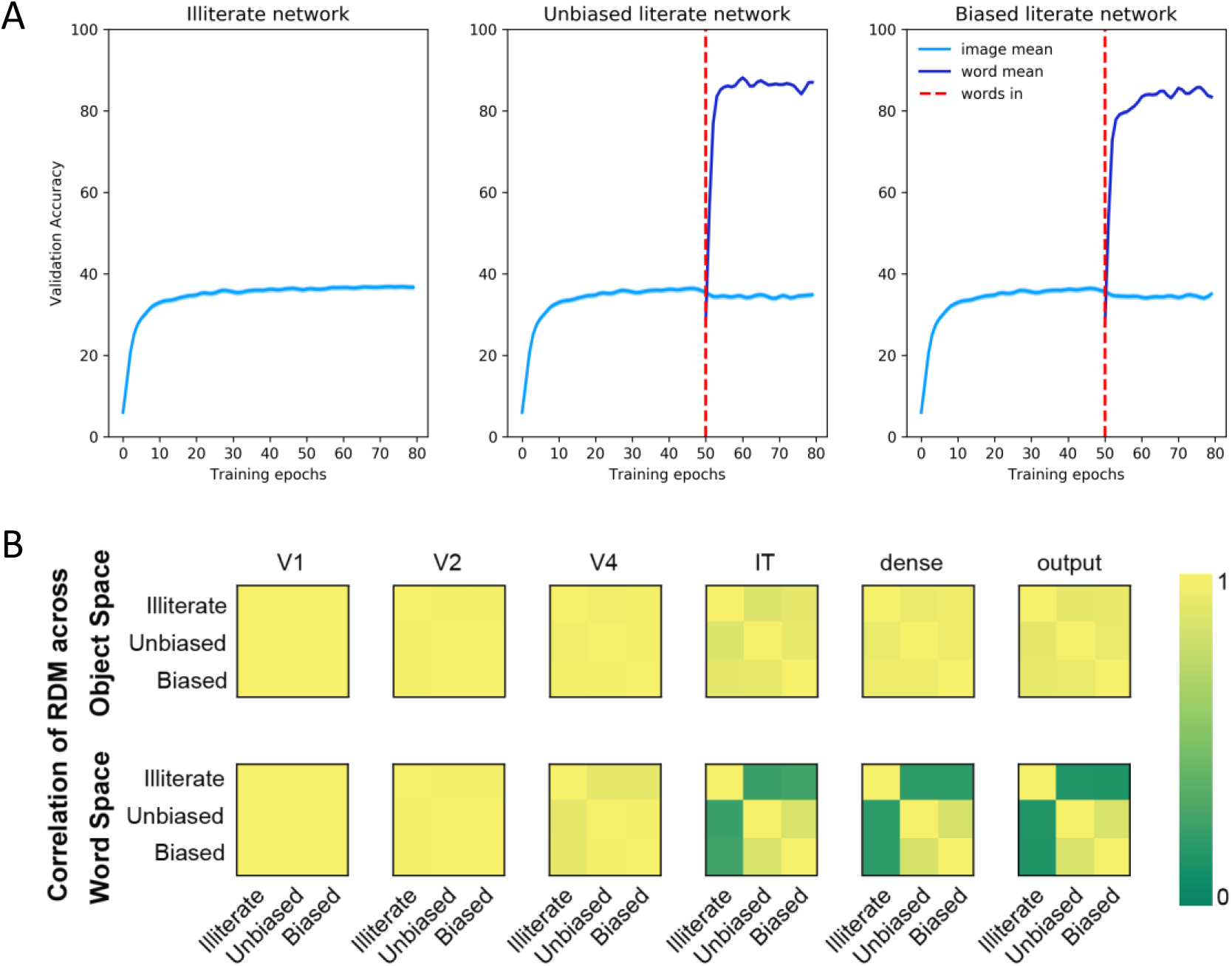
Changes in performance and internal representations during training. A, changes in performance: Average top-1 accuracy in the illiterate, unbiased literate, and biased literate networks on the validation set, separately for images and words. The introduction of words at epoch 50 (marked by a dashed red vertical line) leads to a sudden increase in word recognition performance, with only a very slight decrease for images. B, Evidence for changes in word but not image representational spaces. For each layer (V1, V2, V4, IT, dense and output), the figure shows the pair-wise correlation of the Representation Dissimilarity Matrices (RDM) which characterize the representational space for images (top) and words (bottom). For each layer of a given network, these RDMs estimated the pairwise dissimilarity (Euclidean distance) across 80 images of objects (from ref. [80]) or 80 words stimuli. The correlations between RDMs (color scale at right) indicate that, while image representations were similar in literate and illiterate networks, word representations were changed by literacy. The two literate networks (biased and unbiased) converged to a similar representation of words in all of their top layers, and even slightly in area V4.

Randomly interleaving words with generic image categories during the second phase of training was, in our case, sufficient to avoid interference (even in absence of dropout, which was not used in our simulations). Indeed, in Figure 2A, only a small drop of performance on images is seen upon the introduction of words.

### Changes in representational spaces

To measure whether and how literacy changed the networks’ representations for images and words, we calculated the mean vector of activation evoked by 80 randomly sampled pictures and words within each layer of the models. We then computed their Representation Dissimilarity Matrix(RDM), which characterizes the structure of internal representations, independently of the specific units activated [47,48]. Finally, we examined how the RDMs resembled each other across the three different networks, an approach known as “second-order isomorphism” [49]. As can be seen in Figure 2B, for objects, the RDMs were extremely intercorrelated, with r approaching 1 for V1, V2 and V4 layers, and only a slight drop for IT, dense and output layers. Thus, the image representational space was very similar in all networks and remained largely preserved even after a network is trained for words (mimicking fMRI representational similarity results from ref. [11]). For words, however, the representational changes were dramatic in the late layers of the literate networks. Both biased and unbiased literate networks converge to highly similar representation space (r approaching 0.85), and this representational space was significantly different from that of the illiterate network (r = 0.16 and 0.15 respectively with unbiased and biased networks in the output layer; resampling test across 10000 bootstraps, p<.0005). The change in representational similarity induced by literacy was already detectable in the V4 layer (r = 0.99, 0.91, and 0.91 for correlation between unbiased-biased, biased-illiterate, unbiased-illiterate respectively), but not V1 or V2 – in partial disagreement with fMRI findings that V1-V2, rather than V4, hosts detectable literacy-induced changes in readers of alphabetic languages ([5,17,50]; although see also ref. [51]).

To visualize those internal changes, we plotted the activations evoked by various word stimuli in the subspace of the first three principal component of word-evoked activity, within each of the three top layers (Fig. 3; although this figure shows only one literate network, i.e. unbiased, very similar results were found for the biased network). This plot revealed how the acquisition of literacy led to the emergence of an invariant neural representation of words, with a decrease in the impact of physical parameters (location and size) and an increase in the impact of reading-relevant parameters (word length and word identity). In the illiterate network, activity was primarily segregated by the physical parameters of the words. Retinal location (4 quadrants) was strongly segregated in IT, and remained separated into top and bottom stimuli in the dense and output layers. In the literate networks, word location had a much smaller impact in IT, and became indistinguishable in dense and output layers. A similar change was seen for physical size, which had a massive impact in all layers of the illiterate network, but not the literate ones. Conversely, with literacy, activation became systematically organized according to word length in all three layers. Most importantly, the activation vectors became increasingly segregated by word identity, irrespective of their location and size. Figure 3 (4^th^ column) shows the projection, onto the first three principal components, of the activity evoked by six example words which could be moved rightwards (♦), moved downwards (◂), or scaled in size (★) from the reference image (▪). Only in the literate networks, the responses to a given word were all clustered together, irrespective of those variations, with this invariance increasing from IT to the dense and output layers.

**Fig 3.**
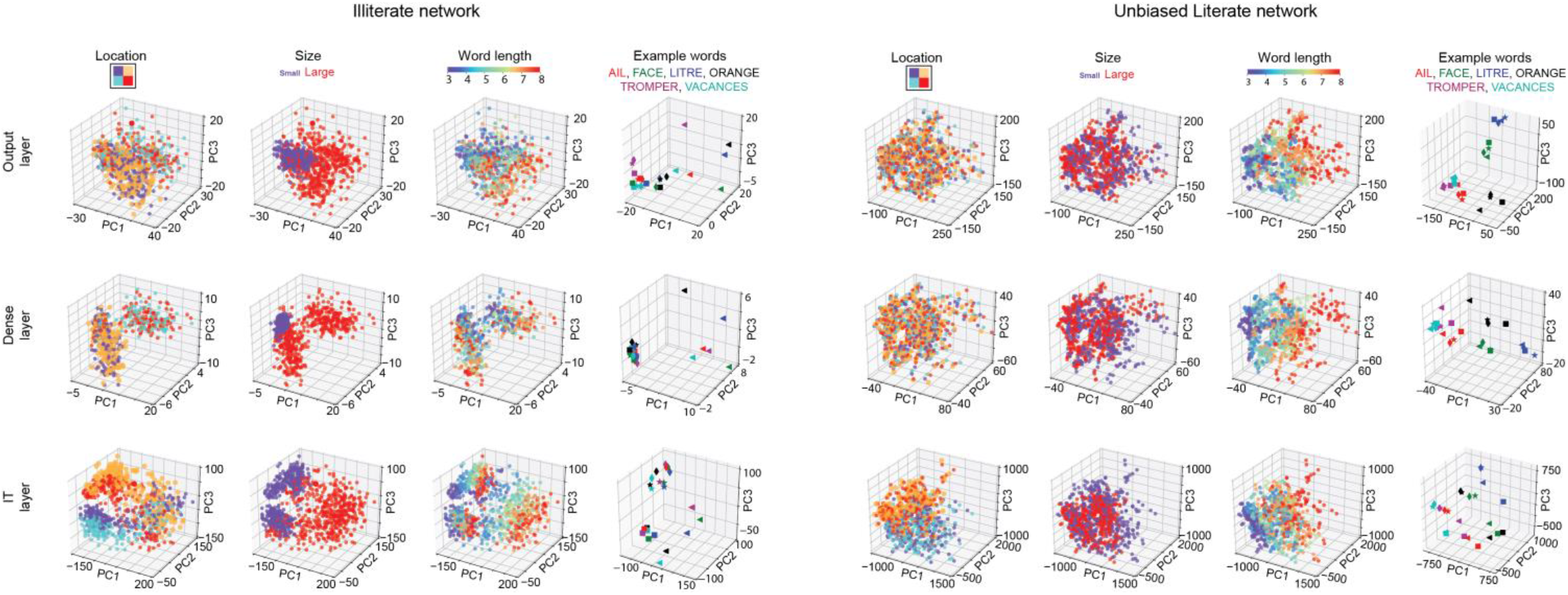
Principal component visualization of word representations. The figure shows the activity evoked by various word stimuli in the top three layers of the illiterate and unbiased literate networks (the biased literate results were similar). Dimension reduction was achieved by extracting the first three principal components in a Principal Component Analysis (PCA) of the activation evoked by 960 stimuli (120 words x4 positions x2 sizes). The location of the words varied from −30 to +30 pixels along the horizontal axis, and −15 to +15 pixels along the vertical axis, thereby spanning all the four quadrants (first column). Their size varied by a factor of two (Arial font at size 40 and 80 points; second column). Their length varied from 3 to 8 characters (third column). The fourth column shows the activity evoked by six example words which could be moved rightwards (♦), moved downwards (◂), or scaled in size (★) from the reference image (▪).

### Single-unit analysis of invariance for size, font and case

We next asked whether we could trace those changes down to the single-unit level. Skilled readers can recognize words effortlessly across changes in size, font, case, and location. At the single-unit level, could we see a trace of those invariances and how they build up across the successive layers? For each unit in the network, and each of three transforms: size, font and case, we collected two vectors of activations. These vectors were 1000-dimensional and recorded the activation evoked in the same unit by each of the 1000 words in the training set, with each word being presented in two distinct randomly chosen values of the transform under consideration (e.g. two different sizes). We discarded changes in location, as location invariance is essentially imposed in convolutional networks by the hard-wired mechanism of weight sharing, but each word location was randomly sampled. We then computed the Pearson correlation coefficient between the two vectors for that transform (see ref. [52] for a related invariance index). The correlation should be close to 1 if and only if the unit exhibits a selectivity profile, across the 1000 words, which does not vary across changes in size, font or case. Repeating this operation for all units within a given network layer yielded a distribution of correlation coefficients (Figure 4A). As a global invariance index for an entire layer, we used the medians of those distribution (Figure 4B).

**Fig 4.**
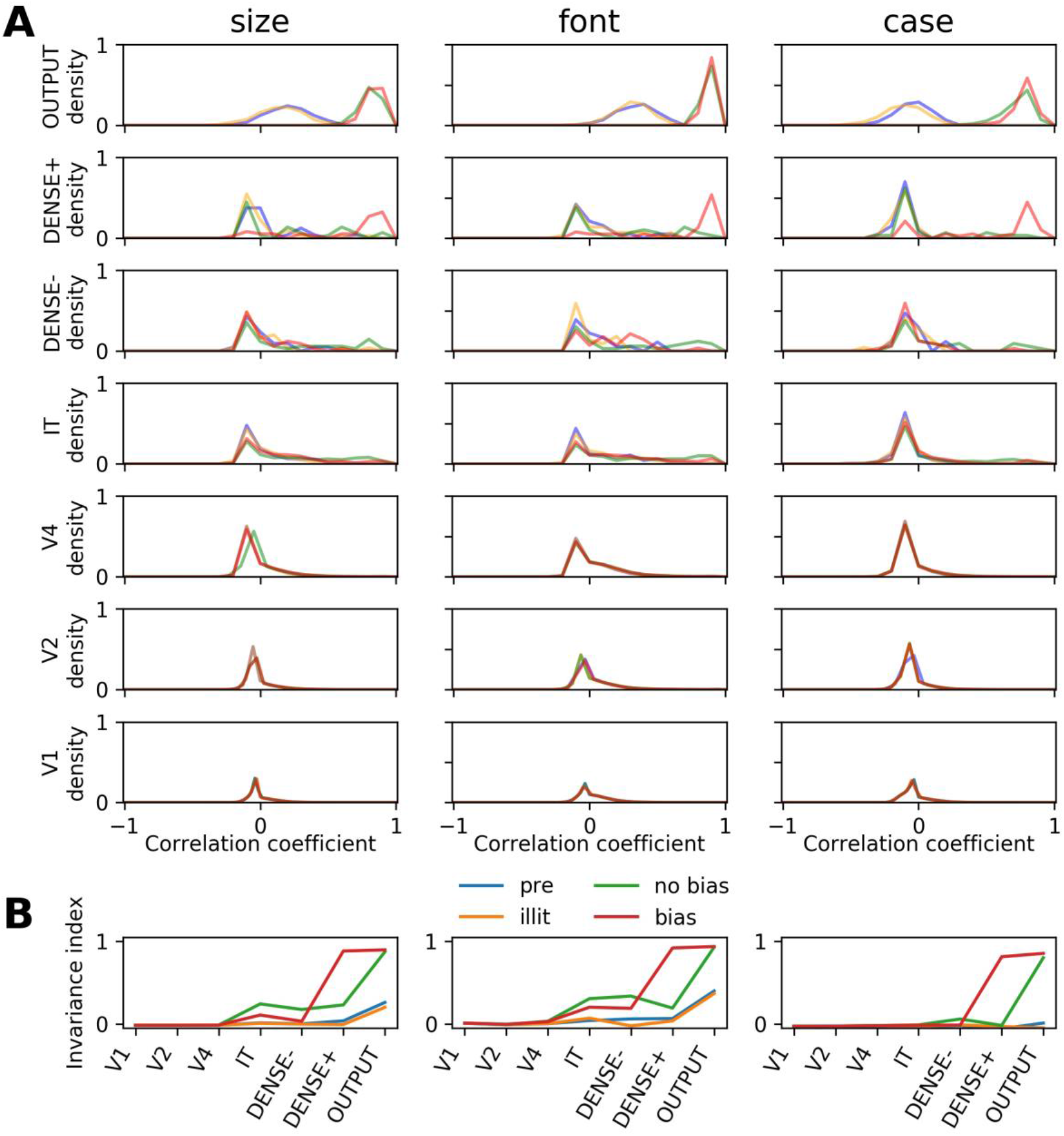
invariance for size, font and case at the single-unit level. (A) Distribution, over all the units of a given layer, of the invariance indices for changes in size, font and case (columns) and for different networks (pre-litterate [pre] in blue, illiterate [illit] in orange, unbiased [no bias] in green, and biased [bias] in red). Each curve shows the distribution, over units in a given layer and network, of the mean correlation coefficient between the activities evoked by two different versions of the word (e.g. font A versus font B). A correlation of zero indicates no invariance. The emergence of a bump near 1 in the top layers of the literate networks (green and red curves) indicates that many units became both word-selective and invariant over irrelevant stimulus variations. Silent units, i.e. units which did not respond to any of the presented stimuli, were excluded from the distribution. (B) Median invariance index(median of the distribution of correlation coefficients) across the hierarchy of layers, for the four different networks.

In the pre-literate (“pre”) and illiterate (“illit”) networks (blue and orange curves in Figure 4), i.e. in the absence of any training with words, the correlation distributions revealed only a modest invariance to word size and font, but not to case. This can be seen in Fig. 4A (top layer), where for both networks, distributions were already significantly shifted towards positive values for size (one − sampled two − sided t − tests performed against zero; 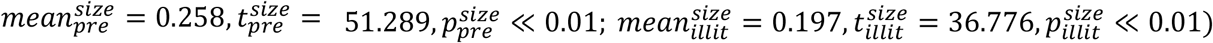 and for font 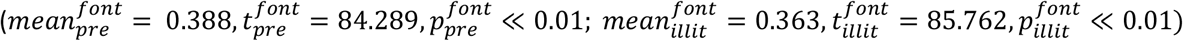 but not different from zero or even shifted towards negative values for case 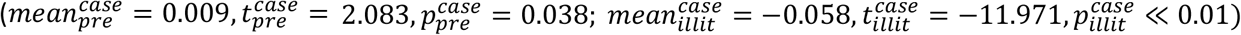. This finding shows that the tolerance to size and to systematic changes in shape that were acquired for the purpose of image classification generalized in part to the novel task of processing words. Contrariwise, letter mappings across upper and lower cases are largely arbitrary (e.g. A and a, E and e), and therefore have to be explicitly learned during reading acquisition [8].

This initial invariance, however, was dramatically enhanced once the networks were trained to recognize words. In both biased and unbiased literate networks (red and green curves in Figure 4), invariance for size and font started to rise in IT, while invariance for case only appeared in the dense layer. For all networks and transforms, invariance always culminated at the output level, where unit responses for a word and its transform became highly invariant (correlation coefficient r close to ∼0.9). Strikingly however, that invariance for all transforms was much stronger in the Dense+ units of the biased network compared to the unbiased network (Figure 4B). This observation shows that biased connectivity promotes the emergence of a restricted cluster of invariant units, mimicking the VWFA [3,6–8], before the final output stage of the network. While this was clearest in the dense layer, the long tail of the distribution for IT units in Figure 4B indicates that many IT units also acquired some degree of invariance, particularly for size and font, in the course of word recognition training.

### Responses to a hierarchy of word-like stimuli

We next examined the responses of the network to a hierarchy of stimuli forming an increasing approximation to real words [29]. The Letter Combination Detector (LCD) model holds that our brains construct visual word representations from raw visual inputs through a hierarchy of increasingly complex and invariant detectors in left vOTC, building up from letter fragments to letters, bigrams, quadrigrams and words [37]. The fMRI study of Vinckier et al. [29] lent support to this hypothesis by showing, in the left vOTC, a posterior-to-anterior gradient of increasing sensitivity to the orthographic similarity of letters strings to real words. However, a recent intracranial study, using the same design but with much higher temporal resolution [31], found this gradient only in the late phase of the response (> 300 ms), where the earlier and presumably bottom-up responses merely distinguished false fonts and infrequent letters from other stimuli. To examine which of those results, if any, were mimicked in the present model, we emulated Vinckier et al.’s study by computing the average activation elicited in each layer of the model by English stimuli mimicking those used by Vinckier et al. (see Supplementary Material, Fig. S1).

All of those word-like stimuli were processed differently in the literate networks compared to the preliterate and illiterate networks (figure 5; table S1). Strikingly, the preliterate and illiterate networks failed to respond strongly to strings, starting from module IT. On the contrary, networks that were trained to recognize words showed increased activations to written words and to word-like stimuli in most layers. Those effects were most prominent in the top layers, and particularly the dense+ units, but were already significant as early as modules V2 and V4. Thus, our simulations mimic the effect of literacy on both high-level visual cortex (VWFA) and low-level visual areas, though not all the way down to V1 [1,11,17,50,51]. Interestingly, in the dense layer of the biased literate network, activation was drastically increased only in the dense+ units that project to word output units, while no change was seen in dense-units (figure 4). This finding shows how the biased-connectivity hypothesis can explain the concentration of literacy-induced changes to a small localized sector of ventral visual cortex [4,5,36,53].

**Fig 5.**
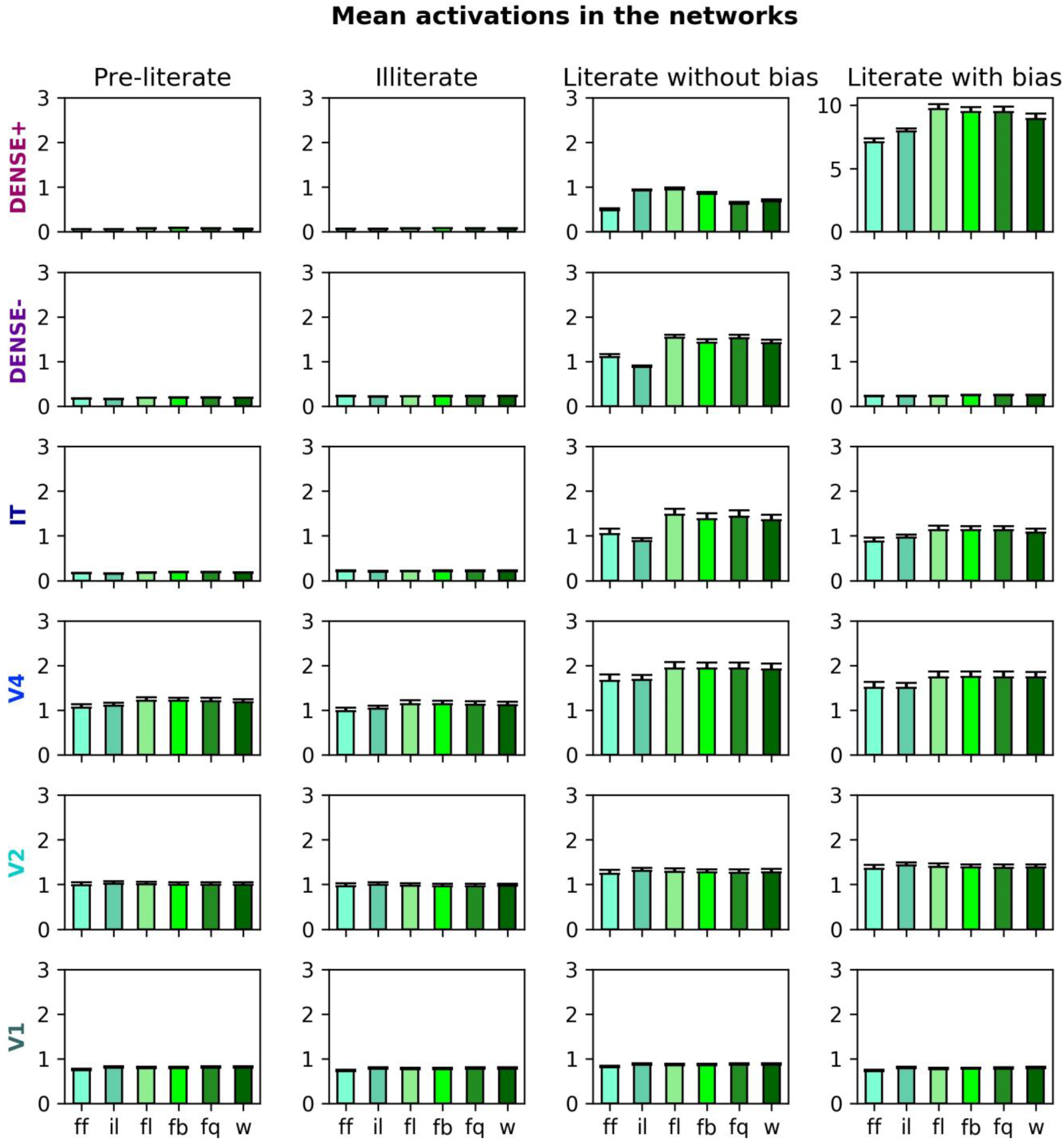
Responses to a hierarchy of increasingly word-like stimuli. Each graph shows the mean activations for each layer and each of the four networks, to a hierarchy of stimuli replicating Vinckier et al.’s (2007) experiment (ff: false fonts, il: infrequent letters, fl: frequent letters, <u>f</u>b: frequent bigrams, fq: frequent quadrigrams, w: words). In their top three layers, networks that were not trained on words showed very little activity to such stimuli, whereas the literate networks showed increased sensitivity, particularly to all stimuli with frequent letters, and particularly in the 49 dense+ units of the biased network (top right graph; notice the different y-axis scale).

While the low activity in illiterate networks implied the absence of strong differences in activation across the 6 categories of increasingly word-like stimuli, such differences were evidenced in the two literate networks, starting in area V4 (table S1). However, those differences were inconsistent with a monotonically increasing gradient of activations across types of stimuli, either in the biased or the unbiased literate networks. Instead of a monotonic gradient, Figure 5 shows a generally lower level of activity for false fonts and low-frequency letters than for all strings that contain frequent letters (frequent letters, bigrams, quadrigrams, and real words). This result is at odds with Vinckier et al’s fMRI study [29], but fits with Woolnough et al’s. intracranial results [31]. With the higher temporal resolution afforded by intracranial recordings, that study was able to distinguish a first bottom-up phase of visual word processing, during which the high-gamma activity was low for stimuli with infrequent letters and soared whenever the stimuli comprised frequent letters; and a second wave (>300 ms) during which the high-gamma activity increased monotonously according to the hierarchy of experimental conditions, as in Vinckier et al. (2007). Since the latter effect started in anterior IT, and then progressed towards the occipital pole, it was tentatively attributed to a top-down effect, with pseudowords closer to lexical items receiving a greater amount of top-down amplification. This conclusion is consistent with the fact that the present neural network simulations, comprising solely bottom-up connections, capture the first effect but not the second. It is also consistent with recent empirical results indicating that the main change induced by literacy is a sensitivity to letter shapes and their precise locations [40,41]. As also concluded by others [35], the simulation of recurrent interactions may be needed to capture the full temporal profile of word-related activations and to mimic fMRI signals.

### Characterization of word-selective units in the dense layer

A typical feature of the VWFA is its activation preference for printed words over other classes of images. Therefore we next asked whether some units in the dense layer of each network exhibited such word selectivity. A unit was deemed word-selective when its average response to word stimuli was 3 standard deviations above its responses to faces, bodies, houses and tools (see Supplementary material for details). As a replication set, we also probed these units’ responses to a hierarchy of increasingly word-like stimuli similar to Vinckier et al. [29].

We found only a few word-selective units in the pre-literate and illiterate networks (resp. 5 and 6 units; Figure 6 top row). Even though those units responded much more strongly to words than to other pictures, their average profile of selectivity did not discriminate between false fonts, infrequent letters, frequent letters, frequent bigrams, frequent quadrigrams and words. We thus interpret these units as being sensitive to the horizontal shape of words and/or to high-contrast line intersections, either as a result of training or by mere chance initialization of network weights.

**Fig 6.**
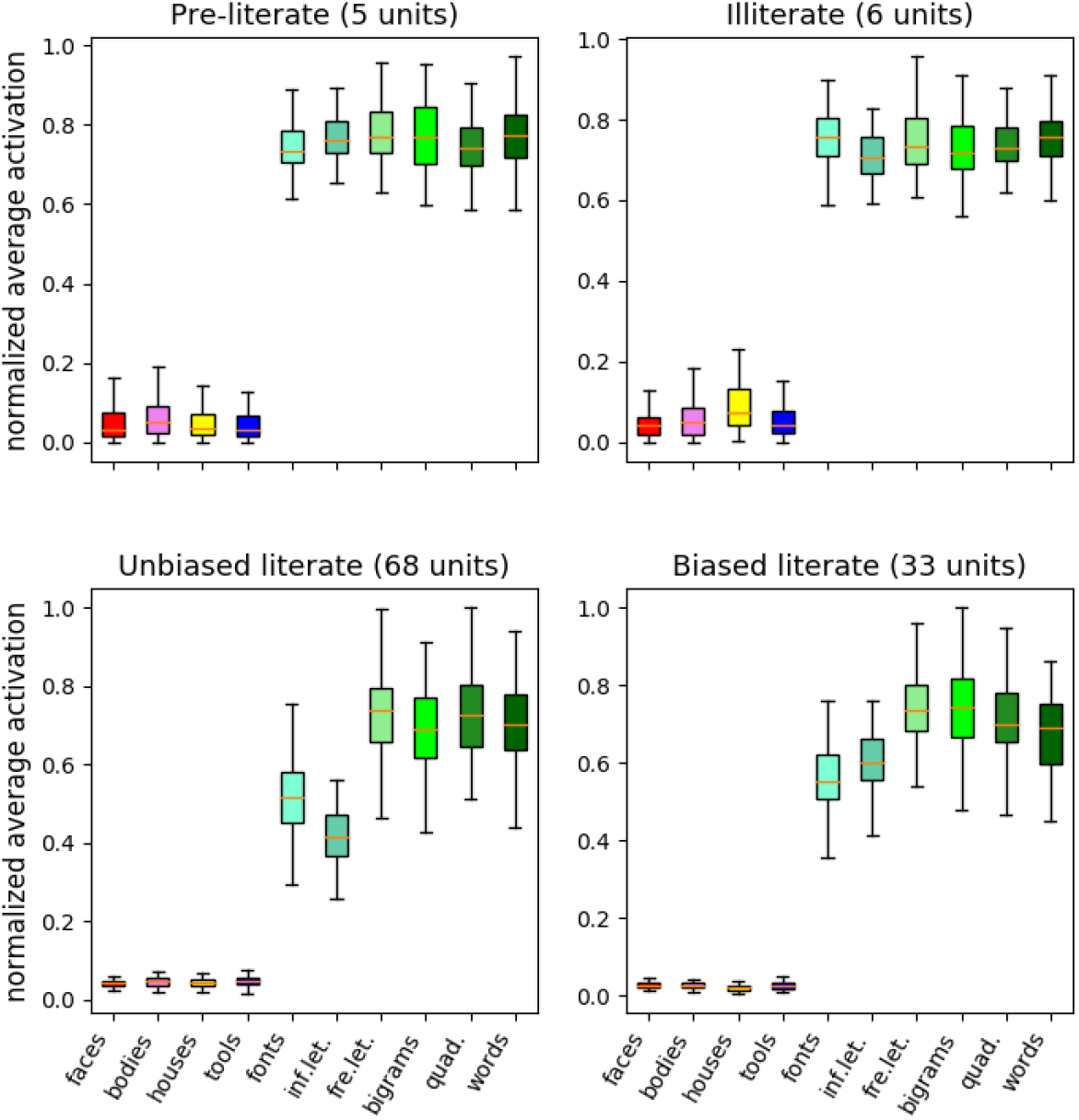
Emerging selectivity for words and word-like stimuli at the single-unit level. Each panel shows the mean activation profile of word-selective units in the four networks, in response to pictures (faces, bodies, houses, tools) and to the Vinckier et al. hierarchy of word and word-like stimuli. Pre-literate and illiterate networks contained very few string selective units, and those did not exhibit any differential sensitivity to letter status or to frequency. Word-selective units were more numerous in the two literate networks, and showed markedly higher responses to stimuli that contain frequent letters.

Word-selective units were much more numerous in the literate networks (33 units in the biased network, including 29 amongst the dense+ units; and 68 units in the unbiased network; Figure 6 bottom row). Those units responded to false fonts and infrequent letters, but showed markedly higher responses to all stimuli that contained frequent letters. This indicates that word-selective units in the dense layer became partially selective for the learned letters, rather than for general word shape or sub-letter features.

We next examined what was the function of those units prior to the acquisition of visual word recognition. Interestingly, we found that prior to the introduction of words in the training set, word-selective units were overwhelmingly uncommitted. In the two literate networks, when returning to the preliterate stage (where the network was solely trained to classify pictures), these units showed no selectivity towards any of the tested categories, and exhibited an overall low response (figure S2). This finding mirrors the longitudinal fMRI findings of Dehaene-Lambertz et al. [11], although in this study, the word-selective units did tend to be initially weakly responsive to tools.

### Effect of word length

We next wondered if the activity of those units would exhibit a length effect, i.e. greater average activity for longer words. The literature on this topic is quite mixed. Behaviorally, for words in a range of about 3 to 8 letters, reading latencies do not increase with word length [54], suggesting parallel processing of letters in the visual system. A significant length effect emerges only for longer words [54], pseudowords [55], words displayed in an unusual format [56], in children before they reach adult reading expertise [57], or in patients with a disrupted VWFA [58], likely reflecting the need for serial processing of letters or other orthographic components. However, reading-related activation in the mesial and ventral occipital cortex, but also in the VWFA, is positively correlated with length irrespective of lexicality and task [31,39,59], suggesting that more neural resources may be engaged in processing longer words even when no serial processing is required. To answer this question in our networks, we computed the activation evoked in word-selective units of the dense layer, for words ranging from 3 to 8 letters. We then evaluated the Spearman rank correlation between length and mean activation.

Figure 7 shows that, on average, the activation of word-selective units in the dense layer of the network increased monotonically with word length. This was true for most units: only a few units show no variation or, very rarely, a negative trend (Figure S3). Such a length effect was restricted to word-selective units in the biased network, while it was stronger but not only confined to word-selective units in the unbiased network. Indeed, the Spearman correlation coefficient was 0.60 in the biased literate network for word selective units, and 0.00 for non-word-selective units. In the unbiased literate network, scores were higher both for word-selective units (0.69) and non-word-selective units (0.15). Thus, the model reproduced the impact of word length on brain activity during word reading (e.g. ref. [31]).

**Fig 7.**
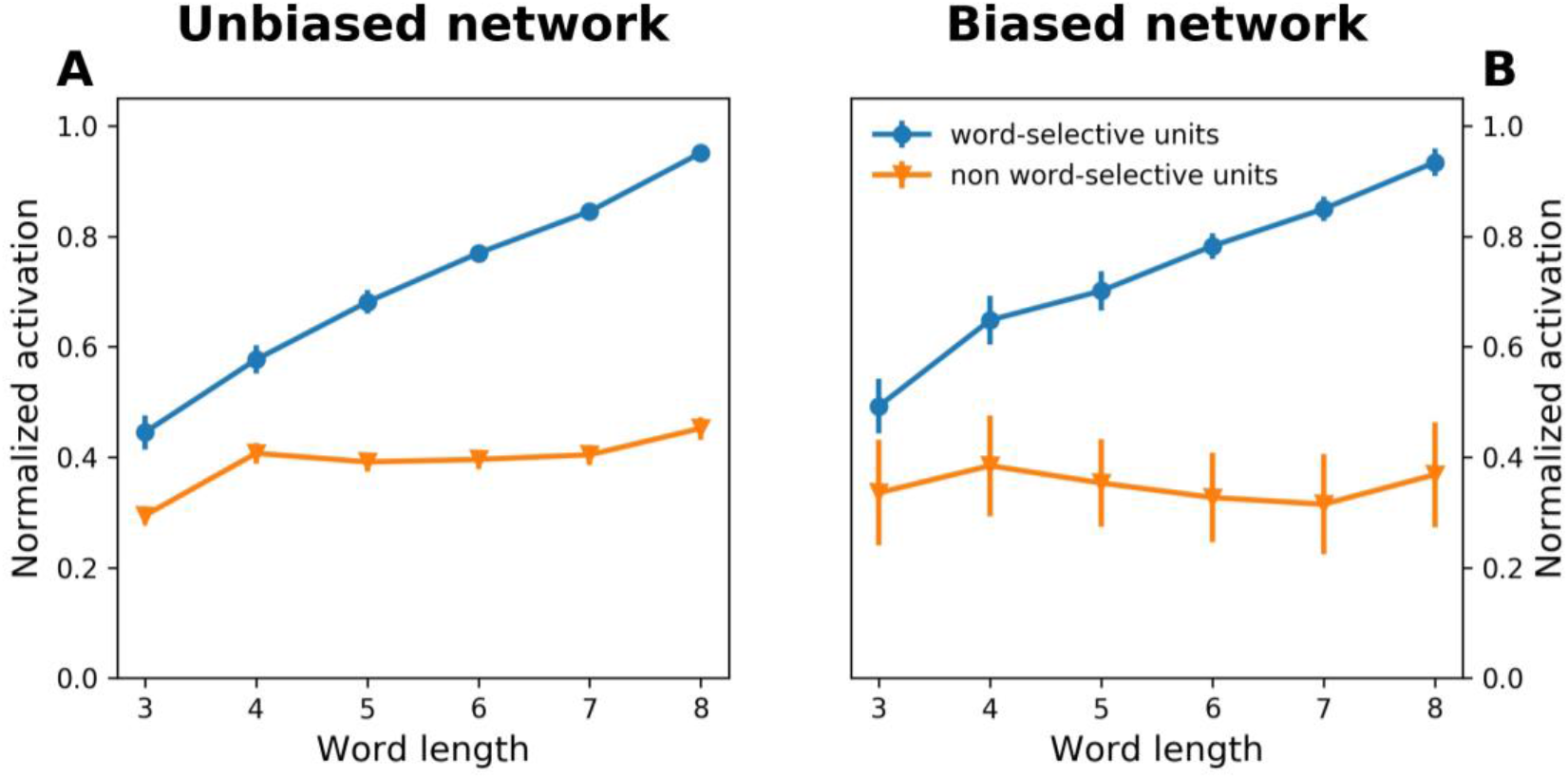
Effect of word length on word-selective units. Activation increased linearly with word length in word-specific units (in both unbiased and biased networks), but remained at a constant low value in non-word specific units.

### Impact of focal lesions: simulating pure alexia

The causal role of the left VOT in reading was recognized in the 19th century from the study of patients with pure alexia [33,34], an acquired impairment characterized by impaired recognition of letters and written words, contrasting with spared general vision, spelling, semantics and phonology. The critical lesion was located more recently to the VWFA as identified with brain imaging methods [34,53,58]. To determine whether this is also the case in the model, we removed different fractions of units in the dense module of the networks, either focusing on biased units (when applicable) or on word-specific units.

Figure 8 (top right) shows the effect of lesions to the dense+ units (or “VWFA units”) in the biased network. For comparison, lesions of equivalent size were performed in the unbiased network (Fig. 8, top left), selecting amongst those same dense units. The performance patterns were strikingly different. The biased literate network became incapable of word recognition even when as few as 20% of its dense+ units were lesioned, while image recognition was barely impacted. The unbiased literate network, on the other hand, barely suffered any impairment in picture or word recognition, even when all 49 dense+ units were removed. Such a selective impairment in visual word recognition is reminiscent of pure alexia, and shows how the biased connectivity hypothesis can account for the existence of a focal site whose lesion selectively affects visual word recognition.

**Fig 8.**
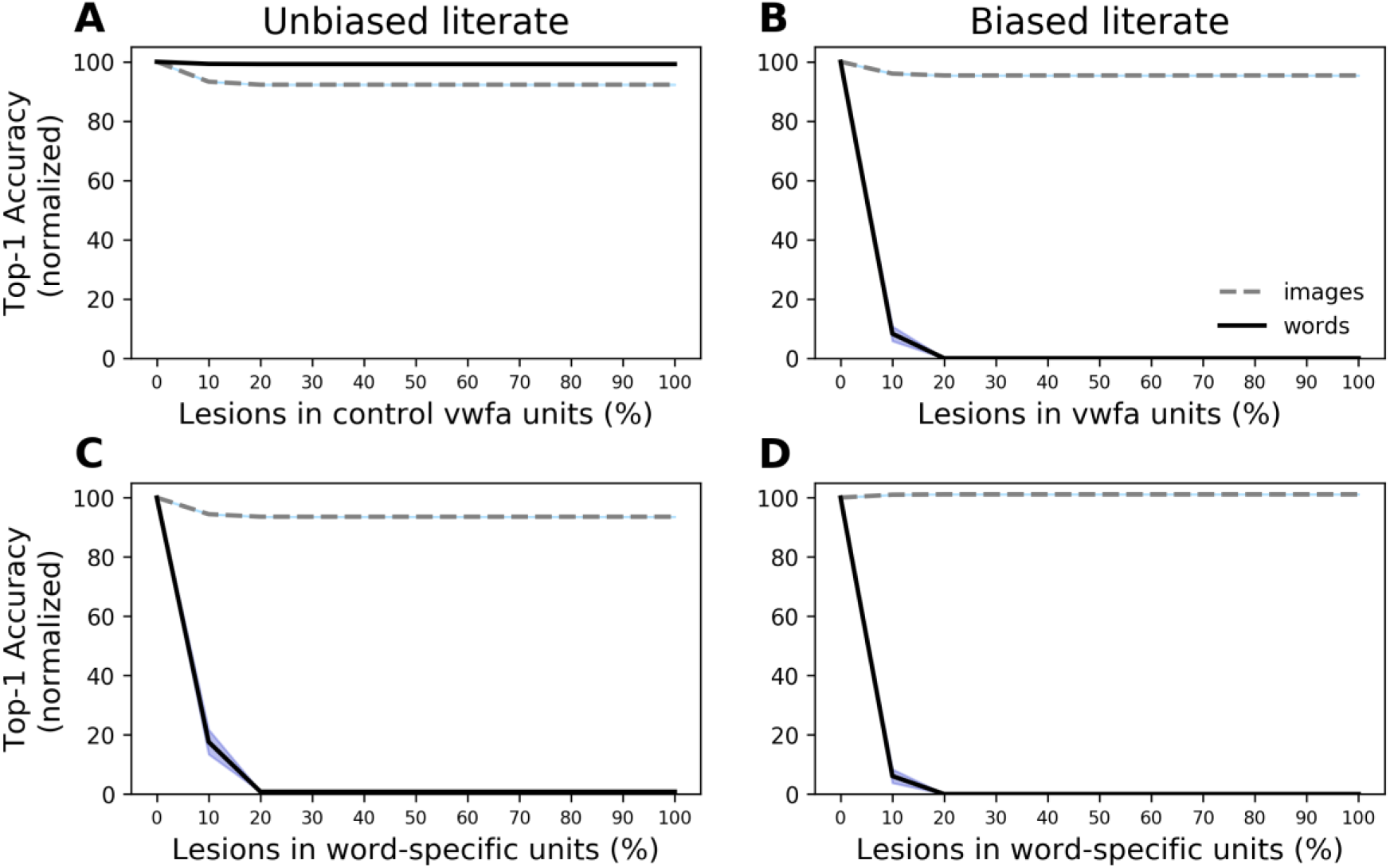
Impact of lesions on picture and work recognition. Classification performance for words and images is plotted against the proportion of units removed, separately for the unbiased (left column) and biased (right column) networks. In the top graphs, the lesions targeted the 49 dense+ units. The top right graphs shows that the biased literate network undergoes a complete loss of performance for words, with preserved image recognition, even when as few as 20% of the dense+ units are lesioned. A similar lesion has very little effect on the unbiased literate network (top left). However, when word-specific units are specifically targeted (bottom graphs), this deficit reminiscent of pure alexia is exhibited by both networks.

Figure 8 (bottom line plots) shows what happens when word-specific units are lesioned: in this case, removing 20% of units in either network suffices to produce a complete impairment on words. This shows that those word-selective units are, indeed, causally responsible for word recognition performance in both networks. However, only the biased-connectivity hypothesis can explain why these units are systematically grouped into a restricted, identifiable cluster that can be targeted by an anatomically cohesive lesion. Also note that with 10% lesions of word-specific units, performance is worse in the biased network than in the unbiased one. This is consistent with the fact that the unbiased literate network has more word-specific units than the biased one and therefore possesses more distributed and more robust neural representations of words.

### A sparse neural code for words

We next analyzed the nature of the neural code that allowed those selective units to encode a thousand words. The first question we asked is to what extent the neural code for words is effectively sparse, using only a subset of active units for a given word, or is distributed across all word-selective units. To this aim, we performed selective silencing experiments. We started by computing, for each unit, its distribution of activity across the 1000 words. Considering the activity pattern produced in all word selective units by a given word, we then set a unit to zero when its activity for this word fell below a threshold percentage of the unit’s global activity distribution, and examined the effect on word recognition performance (solid curve marked “lowest” in Figure 9 A-B). The results showed that it was possible to silence the word-selective units whose response fell in the bottom half of their global distribution without compromising at all the word recognition performance of the networks. Silencing in the opposite order, however, starting with the most responsive units (dashed curve marked “highest” in Figure 9 A-B) had a dramatically different impact. Silencing just the single unit whose response were the strongest resulted in a dramatic drop in performance. This analysis shows that the neural code for words is sparse: not only is it based on a small number of selective units (68 for the unbiased network, 29 for the biased one), but the strongest responses of these units are absolutely necessary for word recognition, while only the top ∼40% strongest responses are sufficient.

**Fig 9.**
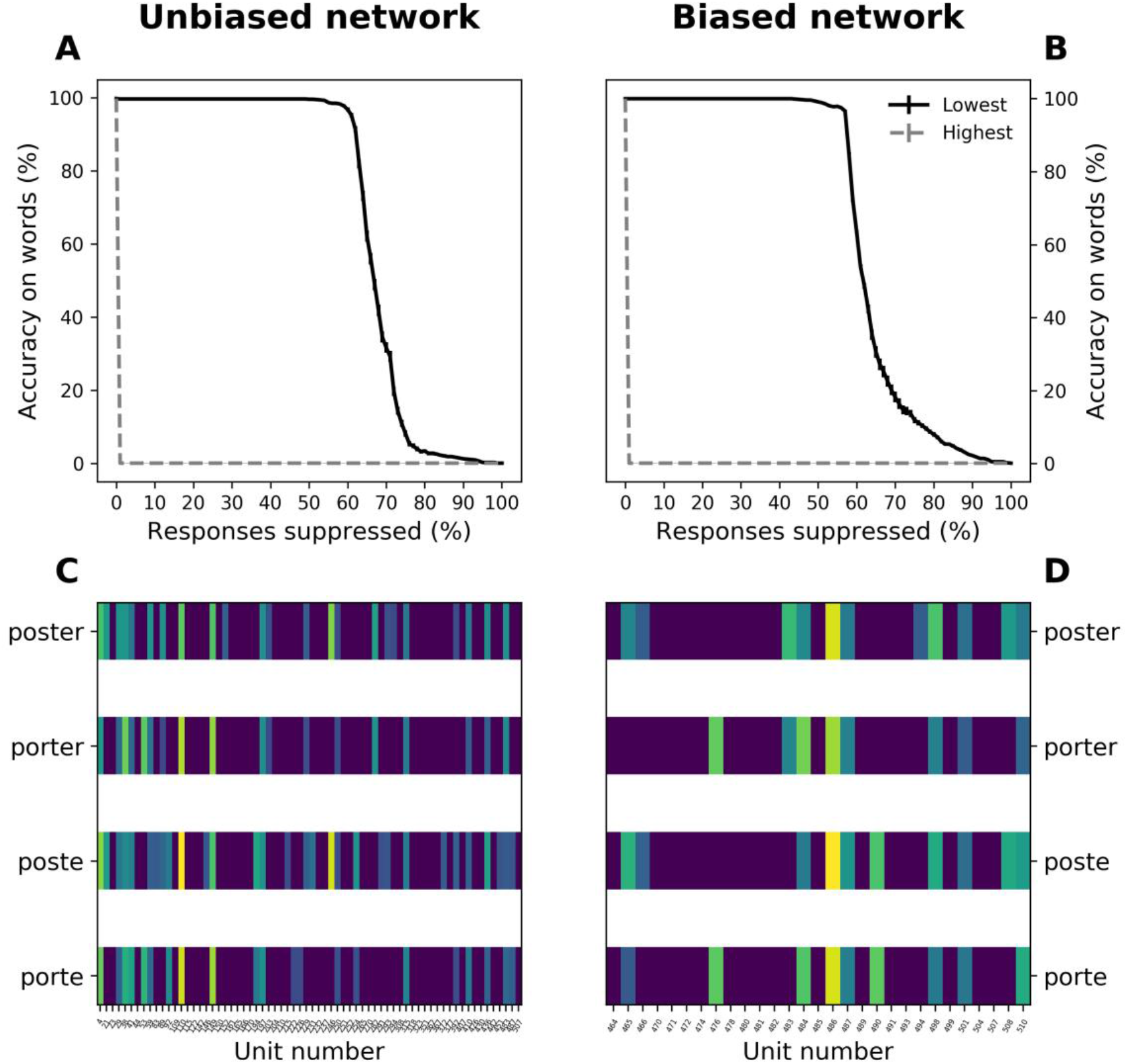
Characterizing the neural code for words. Left column: unbiased network. Right column: biased network. (A, B) Word classification performance as a function of activity cutoffs in word-selective units (normalized by performance score without cutoff). Cancelling the bottom-half responses of each unit leaves classification performance intact (black solid curve), while cancelling the top 1% responses removes all word classification ability (dashed gray curve). (C, D)Example of the sparse neural “barcodes” sufficient to encode a given word, obtained by keeping only the activity above the median of each unit’s activation distribution.

In order to examine the patterns of activity for each word, we silenced the activity of each unit below the median of its activation distribution across all words. This procedure resulted in a sparse “bar code” purified of its unnecessary activity, and yet sufficient to recognize each word (see examples in Figure 9 C-D). Sparsity, defined as the average proportion of units with null activity, was very high (79.3%) for the biased literate network: on average, only 6 out of 29 word-selective units showed non-zero activity to a given word. Sparsity was even higher (86.7%) in the unbiased literate network, with an average of 9 out of 68 units activated by a given word. Thus, even in absence of dropout, which is known to produce sparser codes [46], word representations in the networks were effectively sparse.

### Modelling the receptive fields of word-selective units

A simple hypothesis for the effect of word length is that most units respond to critical features such as letters at a certain location (a bank of letter detectors, which is the front-end in most models of word recognition [60–63]), or bigrams and other letter combinations [37,38,42,64]. Longer words would then have a greater likelihood of activating a greater number of such detectors.

To test this hypothesis, we attempted to model the response of each unit across words as a linear combination of its constituent letters at each of the 8 possible positions (26 letters at each of 8 possible positions). Given the large number of features, we performed cross-validated regularized linear regression (LASSO) to generate a sparse estimate of features that activate a given neuron. The response of each unit was measured along 8000 stimuli, i.e. each of the 1000 words in the training set (3-8 characters long) presented at 4 different locations and 2 different sizes, while the model matrix comprised 26*8 = 208 features. While this number may seem large, note that (1) it is relatively small compared to the 8000 data points of each unit; and (2) the conservative cross-validated Lasso regression attributed null weights to many of those regressors. The LASSO regularization constant was estimated using 5-fold cross-validation.

The letter X position model was able to explain a large and significant proportion of the response variability in word-selective units (Figure 10A). Figure 10C,D shows some of the reconstructed receptive fields. The units with highest letter-model fits were either selective to a single letter irrespective of their position (e.g. unit # 282 responded to S at all positions), or were selective for a single or a few letters at a certain position (e.g. Unit # 23 responded primarily to curved letters at 1^st^ position). Some units were ultra-selective (see Supplementary material, Fig S4), i.e. responsive exclusively to a single letter at a specific location (e.g. unit #142 responded to letter M at the word beginning). Such narrow selectively was not seen inside words, however, where most units were selective for certain letters over a range of nearby positions (e.g. unit #146 responded to letter O anywhere in the middle of the word). Most units were positively activated by certain letters and negatively activated (i.e. inhibited) by others in what could be termed a “letter-dipole” configuration (e.g. unit #33 encoded “E but not R towards the end of the word”). In some cases, the preferred letters occupied distinct locations (e.g. unit #21 seemed to care for letter E at a location left of letter R). In most cases, however, a unit was sensitive (positively or negatively) to letters at roughly the same location, suggesting a factorial code where a product of independent preferences for location and letter determined the unit’s response, as proposed in some theories [65,66] and as observed in neural recordings of monkey inferotemporal cortex [43].

**Fig 10.**
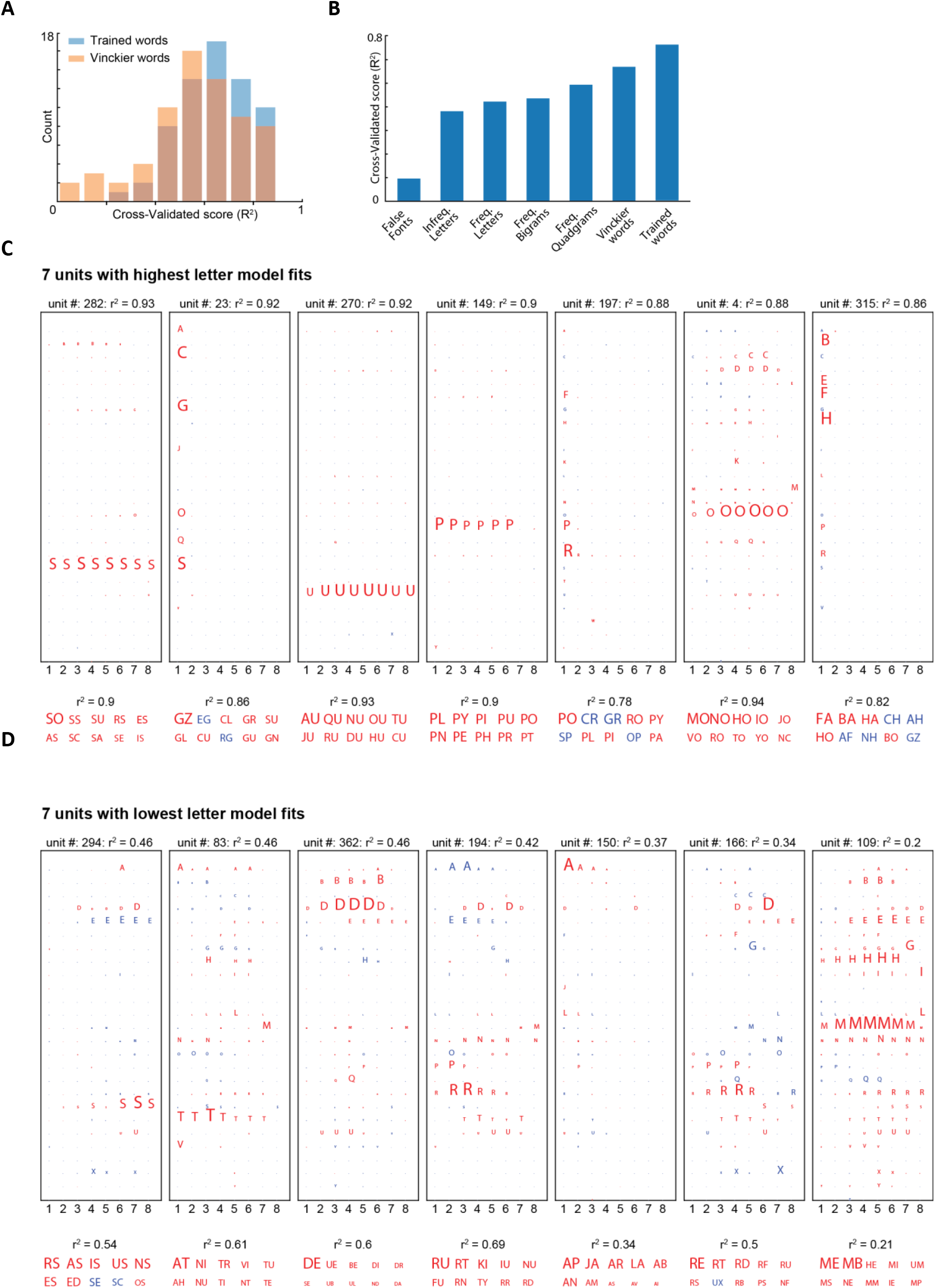
Modeling the receptive fields of each unit. A, distribution, over word-selective units in the dense layer of the unbiased literate network, of the cross-validated model fits (R^2^) from a model which assumes that units respond to a linear combination of specific letters and their location (see text). With 8000 degrees of freedom, all fits were significant, both on the words used to train the linear model and on a new set of words from the Vinckier-like stimulus set. B, Average model fits across different categories of stimuli (false fonts, infrequent letters, frequent letters, frequent bigrams, frequent quadrigrams, Vinckier words, and words used to train the model). C, Top: Examples of reconstructed letter-based receptive fields of the seven word-selective units with the highest letter-model fits, from the non-biased literate network. Each matrix shows, for a given unit, which combinations of letters (vertical dimension, 26 levels) x position (horizontal dimension, 8 levels) were significant predictors of the unit’s response to words. LASSO regression set many coefficients to zero. Non-null coefficients are indicated by the size of the corresponding letter. Positive coefficients are indicated in red, and negative ones in blue. Below: Bigram model fits for the same units, along with the ten bigrams with the highest regression weights. Weight magnitude is indicated by letter size, and sign is indicated by colors (red = positive, blue = negative). D, Same as (C), but for seven units with the lowest letter-model fits.

### Evaluating letter versus bigram codes

We verified that this letter X position model could predict the neural responses to novel stimuli forming a hierarchical stimulus set, i.e. 300 six-letter strings with infrequent letters, frequent letters, frequent bigrams, frequent quadrigrams, and words (see Figure 10B; false fonts were also included as a control for overfitting, since each letter was substituted one-by-one by a roughly similar non-letter shape). Interestingly, we found that the model fit gradually increased from false fonts to word-like stimuli. The increase from high-frequency letters to bigrams, quadrigrams and words contrasts with, but is not incompatible with our previous observation that the *mean* unit activity failed to increase across the hierarchy. It merely suggests that the units became tuned to stimuli that are finely adapted to the statistics of real words. Indeed, Figure 10B is incompatible with the hypothesis that the model units are solely responsive to individual letters: they must also be sensitive to letter combinations.

Indeed, while the proposed letter model was a good overall predictor of the response to words (mean R^2^ = 0.66), the model fits were poor for a few word-selective units, which exhibited a mixed and complex letter X position selectivity (Figure 10D). We wondered if their responses would be better predicted using letter bigrams as regressors. To test this idea, we first modelled the word responses as a linear combination of responses to frequent bigrams. The 1000-word training set comprised 271 unique bigrams (pairs of side-by-side letters, regardless of their position), which were used as features in a second linear regression. Interestingly, the bigram model fits (mean R^2^ = 0.73) were significantly larger than those of the letter model (p < 0.00005, paired t-test). However, the better model fits could be a consequence of the greater number of degrees of freedom of the bigram model. Indeed, according to the corrected Akaike information criterion (AICc), both model fits were comparable and their AICc values did not differ significantly (p = 0.26, paired t-test). For each unit, figure 10CD shows the bigram that had the highest model coefficients. This analysis often confirmed the letter-based model’s conclusions. For instance, unit #282, which responded to letter S at any position, was fitted in the bigram model by approximately equal weights for all frequent bigrams containing that letter (SO, SU, SE, ES, etc). Thus, the most economical description is that this unit simply responds to letter S. However, for units with poor letter-model fits (Figure 10D), the bigram model appeared much more satisfactory, with some units responding strongly to a single bigram (e.g. unit #362, bigram DE) or to two of them (e.g. unit #109, bigrams ME and MB. Thus, some units behaved as approximate bigram detectors, as postulated for instance in the LCD model of word recognition [37].

Next, we asked whether these two models explained distinct variance in the word responses. If true, then a combination of the features from both models (n =26*8 + 271=479) should yield higher fits compared to either of the individual models. This was indeed true: the observed average model fit (R^2^ = 0.77) was significantly higher than either of the two models, and crucially, its average AICc value was also significantly lower (p < 0.00005, paired t-test).

In summary, although the most parsimonious letter-based model explained a large amount of variance across many word-selective units, the bigram model led to equivalent results, and most crucially, the best fit was obtained by combining both regressors. Those findings suggests that, with literacy, the networks became attuned to letters, their absolute position, but although the statistics of their cooccurrence in the trained orthographic system.

## Discussion

Our goal was to develop and assess a minimalist model of how the VOT cortex changes during reading acquisition, through the mere recycling of a biologically plausible convolutional network model of object recognition, without introducing ad hoc reading-specific constraints. As occurs in children, a standard CNN was first trained to identify pictures of various objects and scenes, and then a set of 1000 words of different lengths across variations in location, size, font, and case. Furthermore, we tested the biased-connectivity hypothesis according to which, in literate humans, only part of the VOT cortex becomes specialized for orthographic coding because of its privileged output connections to language areas. To this end, we compared networks whose dense layer was either fully connected to all output units, or in which only a subset of dense units were connected to the output layer, simulating a putative VWFA (Figure 1).

Behaviorally, we found that a network designed for image recognition could easily learn to recognize 1000 written words. Both biased and unbiased networks reached an accuracy of 80% or more after very few training epochs (Figure 2A). Critically, accurate recognition of abstract word identity occurred across large variations in the physical features of stimuli, particularly across letter case, thus attaining the perceptual invariance developed by expert human readers. The performance of biased and unbiased networks was similar, suggesting that the development of an anatomically focal VWFA does not present a functional advantage per se.

The acquisition of reading induced only a barely perceptible deterioration in the previously trained object recognition (note that catastrophic interference [67] was avoided by mixing pictures and words in the training set). This result is compatible with the fact that general object perception abilities differ only minimally between illiterate and literate adults [17,68], and that deficits in object recognition are not typically observed during the acquisition of reading in humans. At the neural level, we found that reading acquisition encroached upon initially uncommitted units that initially exhibited an overall low response to pictures (figure S2). This finding is compatible with longitudinal fMRI data showing that, in the first year of reading acquisition, the VWFA emerges at a site with little or no response to objects or faces [11]. It nuances the “direct competition” or “pruning” view of neuronal recycling [69]: the acquisition of reading probably does not compete with other visual recognition abilities, such as face recognition, by dislodging them from their locations, but by occupying nearby neural sites that cease to be available for the further growth of face-and object-selective regions, thus forcing them to develop elsewhere (for instance in the right hemisphere [11,17]). Here, we concentrated our analyses on the growth of word responses, but in the future, it would be interesting to study whether this growth did indeed compete with the network’s abilities, either to dedicate units to additional pictures such as faces [11,17], or to accurately recognize them across mirror inversions [51,70,71], thus mimicking the small negative downsides of literacy that have been reported in the literature.

At the neural level, we found that reading acquisition led to the structuring of a distinct neural space for words (Figure 3). In its top three layers, corresponding to mid to anterior IT cortex, the network developed critical features of human reading, including invariance for physical location and size, sensitivity to word length, and a segregation of responses according to abstract word identity. Small representational changes were also seen in area V4, but the simulations did not reproduce the reading-related changes in early visual cortex that have been observed in human fMRI when contrasting literate and illiterate subjects [17] or known letters versus control stimuli [5,20,50]. It is possible that those responses reflect top-down inputs from higher-level areas [31,51], which were not simulated in the present, purely feedforward network. However, it is also possible that the convolutional structure of the network, with its automatic duplication of weights across image locations, prevented the emergence of a reading-related retinotopic specialization, as observed in refs. [17,50]. The convolution hypothesis is a massive simplification adopted for computational efficiency, and computational resources prevented us from simulating a non-convolutional network, which might be necessary to capture such fine-grained effects.

At the single-unit level, perceptual invariance emerged from the development, in the top layers of literate networks, of a restricted set of word-selective units whose activation profiles were strongly invariant across changes in size, font and case (figure 4). The most remarkable finding is how compact this representation was. Only 68 word-selective units emerged in the unbiased network, and only 33 in the biased network --and in the latter, only 29 belonged to the dense+ units and could therefore play a causal role in word recognition. Of course, in real-life, each of these dimensions would likely be associated with an entire column of neurons rather a single unit. Still, the results show that a 29-dimensional space (or even less, assuming that further dimension reduction could be achieved) suffices to recognize 1000 words. While surprising, this number is on a par with the 50-dimensional space that demonstrably suffices to encode faces at the neural level [72]. Intuitively, the statistics of letters are highly redundant [73], and conversely a distributed code with 29 dimensions has a large combinatorial capacity (since 2^29^ ≈ 536 million), even when taking into consideration the need for robustness. In fact, we found that, for any given word, the number of required units was even smaller. This is because each word did not evoke a fully distributed code over all word-specific units, but a sparser code: for any given word, silencing the ∼60% of word-selective units most weakly activated by this word did not impair identification. Thus, on average, less than 30 active units (for the unbiased network) or 15 units (for the biased network) sufficed to uniquely specify any of the 1000 trained words.

Figure 10C represents our best attempt to characterize these dimensions of word encoding, i.e. the receptive fields of the word-selective units. We found that most units are sensitive to the presence of one or a small set of letters at a given location, sometimes with a contrast or “letter dipole” (e.g. E but not R). However, we also found that the units’ receptive field could not be solely described by a sum of responses to individual letters: a better fit was achieved by assuming a sensitivity to neighboring letters, i.e. bigrams. The empirical data is conflicting: there is behavioral and brain-imaging evidence that bigrams are a crucial cue to word identity [29,30,37,38,42,74], but also recent data suggesting that the bulk of bottom-up orthographic coding may be based on a conjunction of single letters and their positions [31,40,41]. The present results reconcile both, as they suggest that, in the course of learning, a neural network will make use of all available statistical cues and will develop both letter X position codes and bigram-sensitive units.

With respect to the biased connectivity hypothesis, we found that biased and unbiased networks developed similar representations (e.g. Fig. 2). The main difference was that the biased network developed a more compact representation, with twice fewer word-specific units than in the unbiased network, despite equivalent overall word recognition performance. This compacity came at the expense of a greater sensitivity to focal lesions. By silencing about 20% of the word-specific units, as defined by their category specialization, we made literate networks completely unable to read, thus simulating the main features of “pure alexia”. The unbiased network was more resilient to small lesions, thanks to more diffuse coding over a larger number of units. Indeed, only the biased network could explain how a focal lesion (restricted to the dense+ units, corresponding to the VWFA) could yield a complete loss of reading abilities. In the unbiased network, the word-coding units were dispersed haphazardly, such that it is hard to see how they could be targeted by a single lesion. Nevertheless, we acknowledge that our model does not have a notion of cortical topography. If additional assumptions were added, such that neighboring units tended to respond to similar features, as in Kohonen networks and more recent work [75], then even the unbiased network may end up acquiring a functionally localized VWFA. Thus, the biased connectivity hypothesis is neither falsified nor settled by this study, and adding topography to CNNs [75] would be an interesting future project.

### Limits of the present model

We now address some of the limits of our modelling approach. A first concern is the extreme neurobiological oversimplification of the network architecture we used. Although CNNs may be construed as simplified models of the visual cortex, they lack many properties such as spiking dynamics, neuron subtypes, cortical layers, intricate local connectivity within or between layers and areas, receptor types and densities, temporal delays, realistic learning rules, etc. By construction, the absence of topology in their upper layers also means that none of these models can reproduce the regular topographical organization of category-specific areas in vOTC, and the emergence of the VWFA at a fixed location relative to those functional landmarks [11,17,19]. For the same reason, the left-lateralization of the VWFA falls outside the scope of the model, as it does not distinguish between left and right hemispheres.

These limitations of CNNs as models of the visual cortex are severe, and yet astonishingly, it is now well documented that when trained on a large number of categories, not only can CNNs reach human-level performance on visual classification tasks, but their single-unit responses can also predict neural, fMRI and MEG activation patterns in the visual cortex of human and non-human primates [35,75–80]. Our choice of CNN architectures for modelling vOTC is thus conservative, and since the present models already mimic several known properties of the reading system (e.g. invariance for case, word length effect, pure alexia, etc), we may hope that our networks’ predictions may fit the activity evoked by written words, as well as others did for faces or objects [35,75–80].

Another critique concerns the training set. The ImageNet picture set that we used was never intended to mimic the statistics of the images that a young child is confronted with. Similarly, the printed words we used as stimuli may not mimic the progressive introduction of digits, letters, and words, possibly handwritten, that a child receives in the family and the school. In the future, it may be important to include more realistic training sets (e.g. ref. [77]), although it is unclear how this would change the representations of these categories, and whether the selectivity analyses that we carried out in the present study would come out differently.

The CorNet-Z network that we used is known to provide a good fit to neural activity despite being relatively shallow, entirely feedforward and trained only on ImageNet [28]. Nevertheless, a crucial missing ingredient is the presence of recurrent connections, which are omnipresent in the primate visual system. Several simulations have shown that recurrent neural networks provide better predictors of activity in the human visual system than purely feedforward networks [35,77,81]. Intracranial recordings suggest that recurrent connections may be particularly important in order to simulate the late part of the neural response to words [31], which is likely to contribute massively to fMRI BOLD signals. In this respect, it is worth mentioning that the present model failed to fully capture an important fMRI dataset, by Vinckier et al. [29], who found a progressive and continuous increase in fMRI activity for stimuli approaching the statistics of real words. Our simulations only captured the difference between stimuli with or without frequent letters. It is striking that the predictions fits with the initial, but not the late part of the human intracranial signals evoked by letter strings [31]. In the future, it will be interesting to see if the late appearance of the full Vinckier gradient can be reproduced using a recurrent network architecture.

Finally, the present work was limited to modelling the orthographic component of reading, putatively attached to the ventral visual pathway. Yet, decades of research have shown that reading acquisition involves multiple routes and a complex set of visual, phonological, morphological and lexico-semantic representations [16,60,82,83]. Other modelers have made the choice of capturing these multiple routes in neural network models, often with great success [e.g. 60–64]. Such global modelling of reading, however, usually come at the expense of not accurately modelling the early visual stages of word recognition; indeed, in those models, the input is usually a bank of abstract letter identities rather than an actual image of the word (for exceptions, see refs. [84,85]). Our networks face the converse limitation: they only process visual inputs up to the classification level, akin to lexical access, without any influence from downstream phonological, morphological or semantic systems. Fortunately, evidence from baboons and monkeys, who have none of those late stages, suggests that visual demands alone may suffice to drive the emergence of orthographic abilities and a VWFA somewhat analogous to humans [43,86,87]. Nevertheless, focusing on the input orthographic system is a design choice for a first modelling step, and not a theoretical statement, as it is known that phonological representations, for instance, do impact on ventral visual representations in humans [39]. Again, a more complex, recurrent architecture, combining visual and phonological inputs, would be needed to accurately capture those observations.

### Conclusion and summary of main predictions

Beyond reproducing existing data, the main purpose of our approach was to develop predictions concerning the fine-grained neural code for words, which may soon become testable either with very high resolution high-field fMRI or with high-density intracranial recordings. We therefore end by summarizing the predictions that arise from our simulations.

First, we predict that the readers’ inferotemporal cortex contains a sparse neural code for written words, with only ∼30 dimensions of vector coding sufficing to encode the 1000 most frequent words. Second, we predict that those vector codes are encoded by word-specific neurons that do not respond to other pictures (faces, bodies, houses or tools). Third, these neurons should be initially uncommitted in the young pre-literate brain, and only become attuned to letters and their combinations in the course of reading acquisition. Fourth, their overall activation should increase as a function of word length. Fifth, the receptive field of those units should be characterized by a high selectivity for one or a few letters at one or several consecutive locations in the string, with occasional sensitivity to order (a neuron may be selective to letter x around position p1 and to letter y around position p2) and/or an additional inhibition to several other letters (thus forming a “letter dipole” responding to letter x but not letter y around position p).

Finally, according to the biased-connectivity hypothesis, the location of those units should be predictable from their pre-existing pattern of projective connections to downstream language areas. And, since we found that even in the biased literate condition, not all dense+ units ended up being selective to words, we predict that within the human VWFA, unlike monkey face patches [88], some neurons may not respond to words and remain committed to other visual categories. The latter predictions, however, may have be revised once more realistic, topographic and recurrent models of reading acquisition are developed.

## Supplementary materials

### Model

All networks were trained on a cross-entropy loss by Stochastic Gradient Descent, with an initial learning rate of 0.01, scheduled to be divided every 10 epochs by a factor of 10, a momentum of 0.9, a weight-decay of 0.0001 and a batch-size of 100, without batch-normalization and without dropout.

A full description of the CorNetZ network architecture is available in ref. [1]. The python script for all network architectures used in this article, and the script for training these networks can be found at https://github.com/THANNAGA/Origins-of-VWFA, along with pretrained network files for all epochs.

### Evaluation of performance

Top-1 accuracy was calculated at every epoch for all 50 validation exemplars in each category. The curves in Figure 2 (main text) present the average and the standard error of the mean across the 1000 Top-1 scores for images (light blue curves) and for words (dark blue curves).

### Principal component analysis

To create figure 3 in the main text, we performed a Principal Component Analysis (PCA) of the activation evoked by 960 stimuli (120 words x4 positions x2 sizes) in last three layers (IT, Dense, and Output) in each of the three networks. The location of the words varied from −30 to +30 pixels along the horizontal axis, and −15 to +15 pixels along the vertical axis, thereby spanning all the four quadrants (first column). The length of these words varied from 3 to 8 characters (third column), and were presented using Arial font at size 40 and 80 points (second column).

### Average network activation across levels

The selectivity analyses presented in Figure 5 (main text) were conducted using stimuli with the same properties as in the original study of Vinckier et al. [2]. Their design included 5 types of letters strings, differing based on the frequency of their component letters, bigrams and quadrigrams in the written French language, plus strings of pseudo-letters (“false font”). The 5 types of letters strings were: (i) strings in which letters, bigrams, and quadrigrams were infrequent (“infrequent letters”), (ii) strings in which letters were frequent but bigrams and quadrigrams infrequent (“frequent letters”), strings I which letters and bigrams were frequent but quadrigram infrequent (“frequent bigrams”), strings in which all components were frequent (“frequent quadrigrams”), and (v) real words. We designed our test stimuli so as to achieve the same hierarchy as in Vinckier et al., but based on the letter combination statistics of the word training set.

**Fig. S1.**
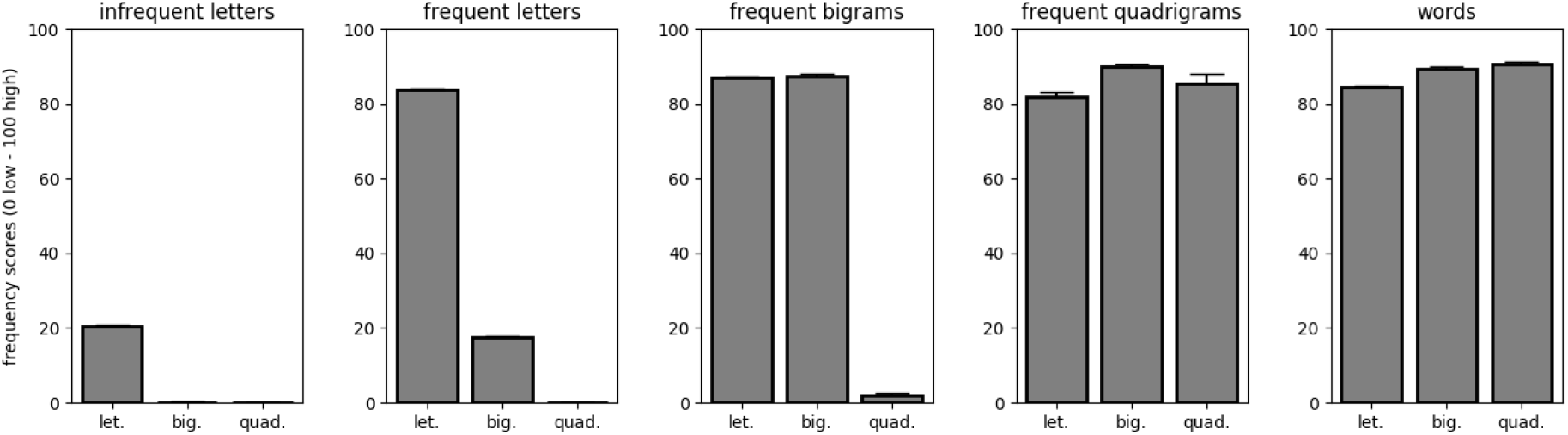
Distribution of frequencies for the stimuli used to replicate Vinckier et al.’s experiment [2].

Each stimulus category is composed of 40 strings. We compute the frequency scores for letters, bigrams and quadrigrams in the following way. Consider a substring **s** of length **ls** from an arbitrary string, and the list of length **l** sorted by descending frequencies of all substrings of size **ls** that were seen during training. The **frequency score** of substring s is defined as zero if **s** does not appear in the list, and as **100*(1 -rank(s)/l)** otherwise. Fig. S1 shows the average frequency score of letters, bigrams and quadrigrams for the 5b classes of our real-letter stimuli, nicely paralleling the Vinckier hierarchy. The false font condition from Vinckier et al. [2]The false fonts used in our simulations were the same as that used in Vinckier et al. [2].

For each category of false fonts (ff), infrequent letters (if), frequent letters (fl), frequent bigrams (fb), frequent quadrigrams (fq) and words (w), 100 uppercase, centered string exemplars were presented to the networks. For each stimulus, in each network module, we then flattened and concatenated the activation values produced by this exemplar in all feature maps of the module, before averaging across all units, hence obtaining an average module activation for a given exemplar. Figure 5 (main text) presents these module activations, further averaged across all exemplars of a given category.

For each network and each module, we carried out 3 one-way Anovas across stimuli categories, testing whether their averages differed for different categories (Table S1). The first Anova (Anova_1to6 hereafter) tested for the difference in means between the 6 categories of stimuli. The second Anova (Anova_1to2 hereafter) tested for the difference in means between the false font and infrequent letters categories of stimuli. Finally, the third Anova (Anova_3to6 hereafter) tested for the difference in means between the most word-like categories of stimuli (frequent letters, frequent bigrams, frequent quadrigrams and words). Table S1 shows the results:

**Table S1.** Results of 3 one-way Anovas, testing in each network and each module, for equal average activations across categories of stimuli from Vinckier et al. (2006). Anova_1to6 tested for the difference in means between all 6 categories of stimuli. Anova_1to2 tested for the difference in means between the false font and infrequent letters categories of stimuli. Anova_3to6 tested for the difference in means between the 4 most word like categories of stimuli, involving frequent letters.

|  |  | **Pre-literate** | **Illiterate** | **Unbiased literate** | **Biased literate** |
| --- | --- | --- | --- | --- | --- |
| **Dense+** | Anova_1to6 | F=27.86, p<0.0001 | F=5.4, p<0.0001 | F=96.09, p<0.0001 | F=153.28, p<0.0001 |
|  | Anova_1to2 | F=1.75, p=0.187 | F=0.64, p=0.425 | F=404.43, p<0.0001 | F=83.01, p<0.0001 |
|  | Anova_3to6 | F=16.99, p<0.0001 | F=1.26, p=0.286 | F=68.64, p<0.0001 | F=18.77, p<0.0001 |
| **Dense-** | Anova_1to6 | F=69.52, p<0.0001 | F=3.16, p=0.008 | F=444.33, p<0.0001 | F=17.25, p<0.0001 |
|  | Anova_1to2 | F=30.56, p<0.0001 | F=2.24, p=0.135 | F=211.88, p<0.0001 | F=1.2, p=0.274 |
|  | Anova_3to6 | F=3.23, p=0.022 | F=1.92, p=0.124 | F=20.13, p<0.0001 | F=13.55, p<0.0001 |
| **IT** | Anova_1to6 | F=67.45, p<0.0001 | F=3.53, p=0.003 | F=419.98, p<0.0001 | F=143.97, p<0.0001 |
|  | Anova_1to2 | F=25.02, p<0.0001 | F=1.7, p=0.192 | F=123.89, p<0.0001 | F=65.76, p<0.0001 |
|  | Anova_3to6 | F=3.95, p=0.008 | F=2.02, p=0.109 | F=22.32, p<0.0001 | F=12.89, p<0.0001 |
| **V4** | Anova_1to6 | F=24.68, p<0.0001 | F=19.89, p<0.0001 | F=57.61, p<0.0001 | F=73.72, p<0.0001 |
|  | Anova_1to2 | F=8.53, p=0.004 | F=8.31, p=0.004 | F=0.8, p=0.371 | F=0.15, p=0.7 |
|  | Anova_3to6 | F=1.68, p=0.17 | F=0.68, p=0.562 | F=0.43, p=0.734 | F=0.14, p=0.933 |
| **V2** | Anova_1to6 | F=1.48, p=0.193 | F=2.42, p=0.034 | F=2.01, p=0.074 | F=3.11, p=0.008 |
|  | Anova_1to2 | F=5.91, p=0.015 | F=7.77, p=0.005 | F=7.93, p=0.005 | F=10.76, p=0.001 |
|  | Anova_3to6 | F=0.2, p=0.898 | F=0.29, p=0.831 | F=0.34, p=0.797 | F=0.34, p=0.794 |
| **V1** | Anova_1to6 | F=7.42, p<0.0001 | F=7.38, p<0.0001 | F=7.3, p<0.0001 | F=7.34, p<0.0001 |
|  | Anova_1to2 | F=36.3, p<0.0001 | F=35.8, p<0.0001 | F=35.56, p<0.0001 | F=35.39, p<0.0001 |
|  | Anova_3to6 | F=1.05, p=0.37 | F=1.04, p=0.372 | F=1.02, p=0.382 | F=1.03, p=0.38 |

### Characterization of word-selective units in the dense layer

To probe category selectivity at the single-unit level, we selected stimuli in the faces, houses, bodies and tools, similar to those used in classical fMRI experiments (e.g. refs. [3]). ImageNet comprised two classes that were close enough to bodies and houses (resp. “Bikini” and “Boat House”). For tools, we used the “ALET” tools dataset, which contained 2345 images sorted into 53 tool classes of at least 200 instances each. For faces, we used the “Caltech Faces 1999” dataset which contained 450 images of 896 x592 pixels, with 27 unique people under different lighting/expressions/backgrounds.

For the analyses presented in Figure 6, we first determined the selectivity of each unit in the dense layer of each network. A unit was deemed selective for a given category if its average responses to exemplars of this category was three standard deviations above its average responses for exemplars of all other categories (i.e. words > faces, bodies, houses, tools). A unit was deemed uncommitted (i.e. “labile”) when it was not selective for any of the tested categories. We also examined the mean response of each unit to the above Vinckier et al. [2] word and word-like stimuli. Finally, in order to investigate the initial tuning of these word-selective units, we conducted the same analysis at epoch 49, prior the introduction of words. Figure S2 shows the selectivity at epoch 49 of units that ended up being word-selective at the end of training. There were 6 such units in the illiterate network, 68 in the unbiased literate network and 33 in the biased literate network (note that of these 33 units, 29 are located in the biased region, while 4 lie outside of it, thus being unable to contribute to word recognition). It is clear that in all 3 networks, prior to the introduction of words in the training set, word-selective units are overwhelmingly uncommitted to any of the categories tested for, mimicking the longitudinal fMRI study of reading acquisition [4].

**Fig. S2.**
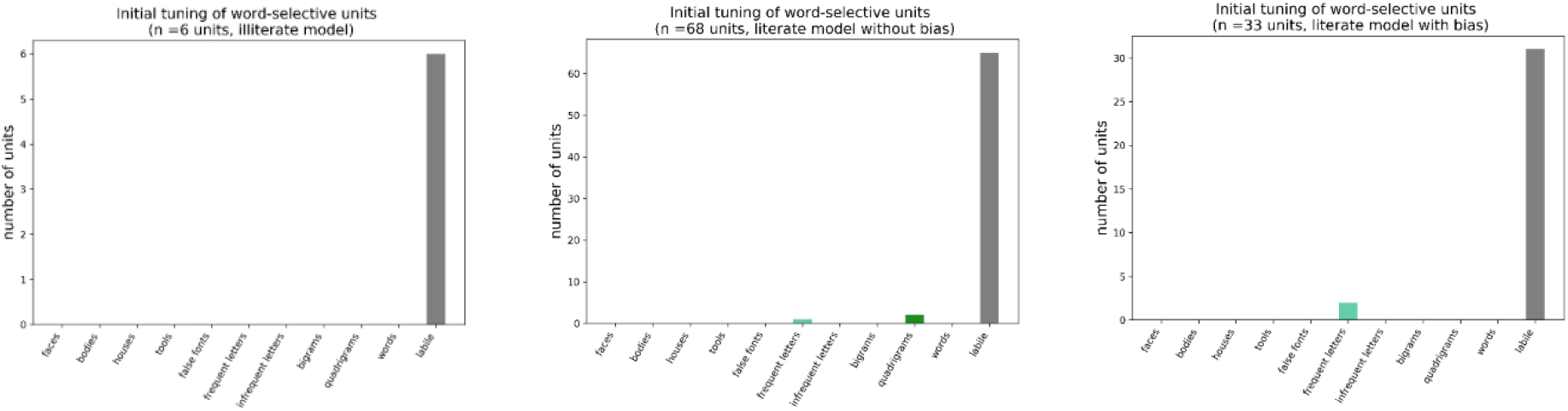
Original tuning of word-selective units in the illiterate network (left), unbiased literate network (middle) and biased literate network (right). In all conditions, word-selective units overwhelmingly come from labile units (i.e. uncommitted units, which were not selective to any of the tested categories).

### Lesion analysis

The upper line of Figure 6 (main text) presents lesion analyses in the unbiased and biased network, focusing on dense+ units. In the unbiased network, lesions were carried out within the same pool of 49 units as in the biased network (units that had the same indices in the dense layer). This allowed a fair comparison with lesions of the same size carried out in biased and unbiased networks. The bottom line of Figure 6 (main text) shows the effect of focusing the lesions on word-specific units across the dense layer. Since the unbiased and biased networks have different absolute numbers of word-specific units that can contribute to word classification (respectively 68 vs 29), for an equal percentage of units (x axis in figure 6), more units end-up being lesioned in the unbiased network.

Performance was calculated as Top-1 accuracy over all ImageNet and word classes in the validation set, using one exemplar per class and averaging, for each proportion of lesions, over 50 random samples of lesioned units.

### Effect of length

Figure 8E and 8F (main text) summarize length effects in word-specific units of the unbiased and biased networks. In Figure S3 below, we present average activations over the 1000 words in the training set, for each unit in both networks as a function of word length.

**Fig. S3.**
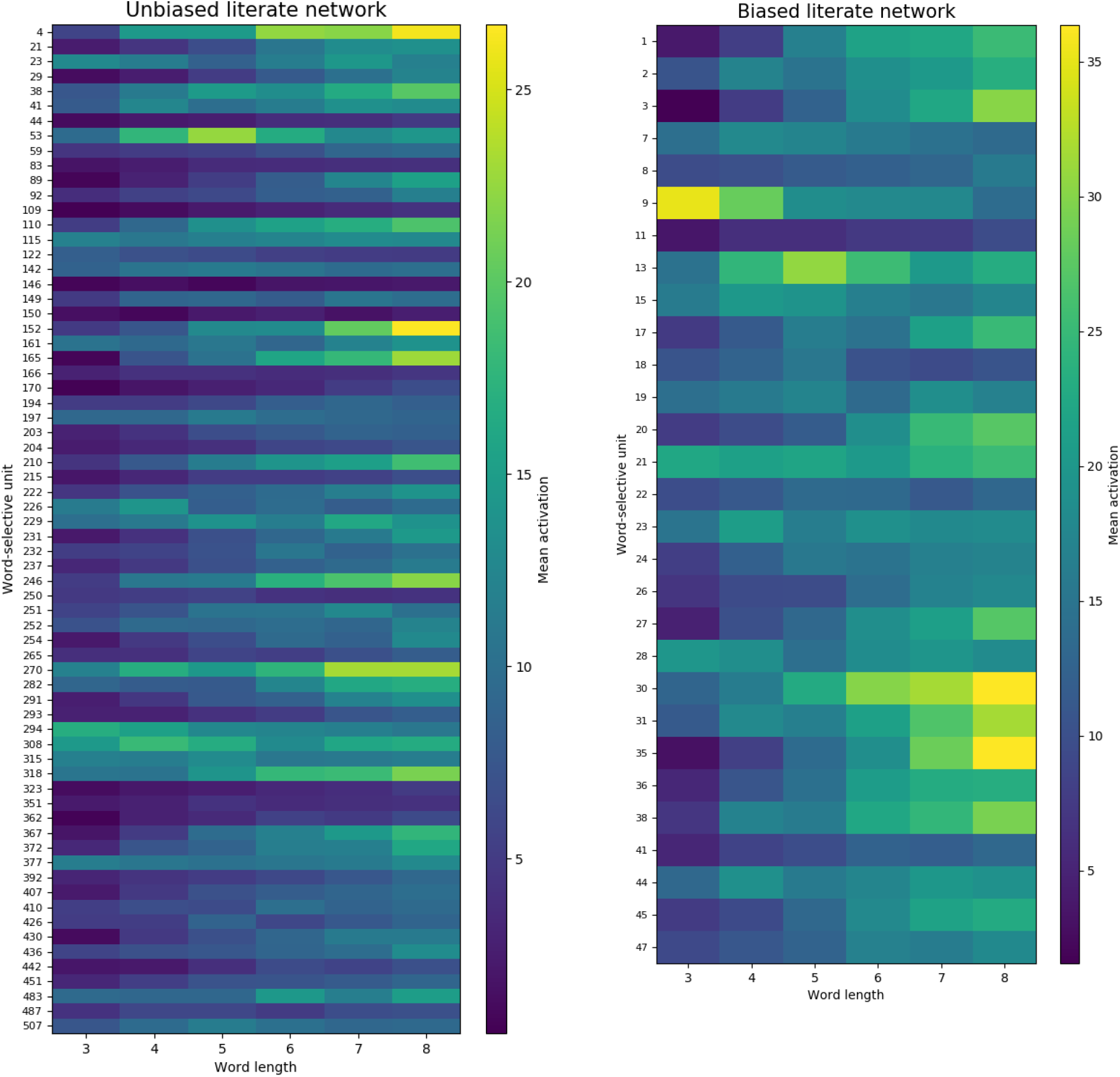
Average activation of each of the word-selective units in the literate networks for words of lengths 3 to 8.

### Visualizing the letter model coefficients

For completeness, figure S4 shows the reconstructed receptive fields of all the word selective units in the unbiased literate network, by visualizing the model coefficients for each letter and each position:

**Figure S4:**
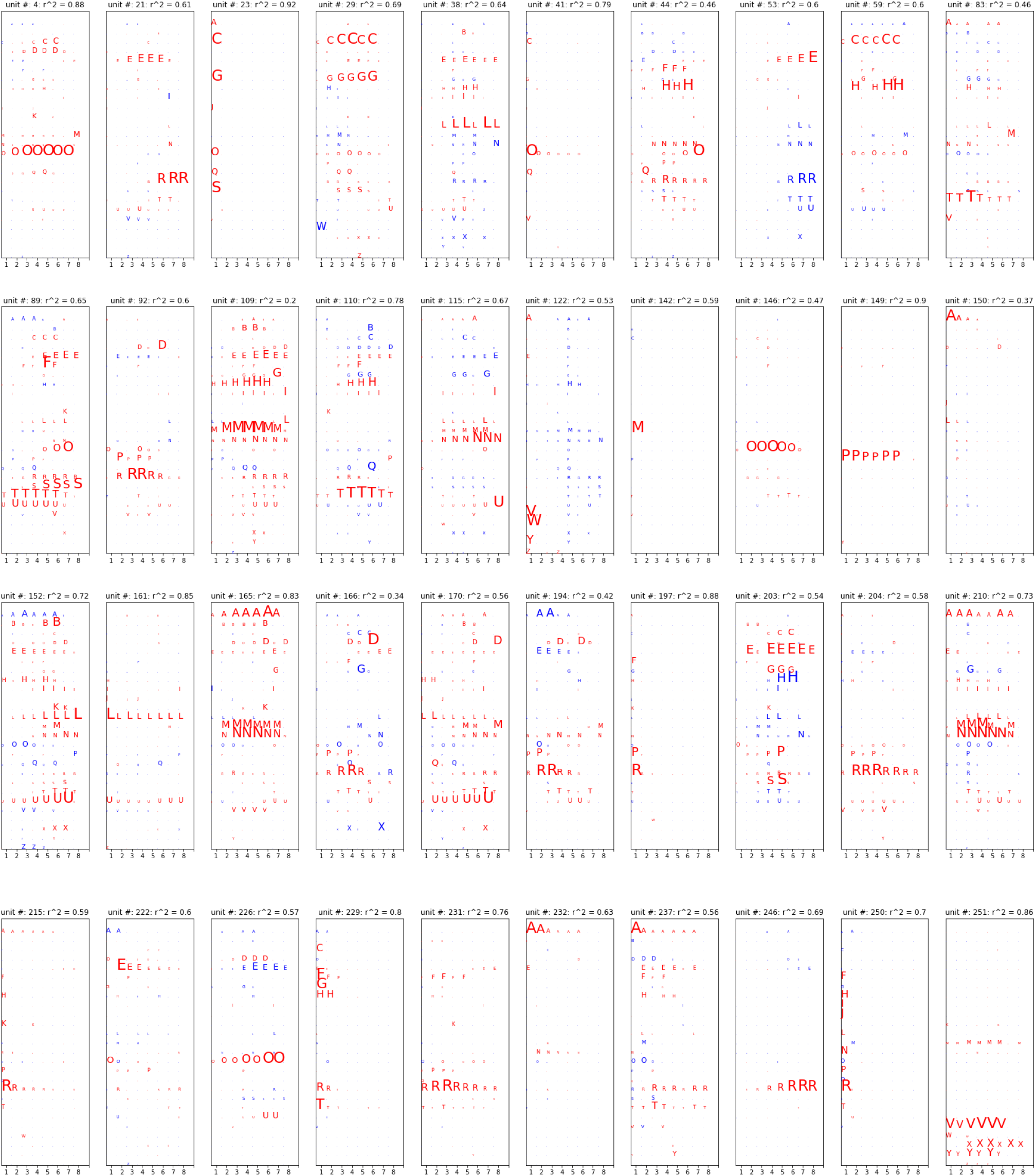
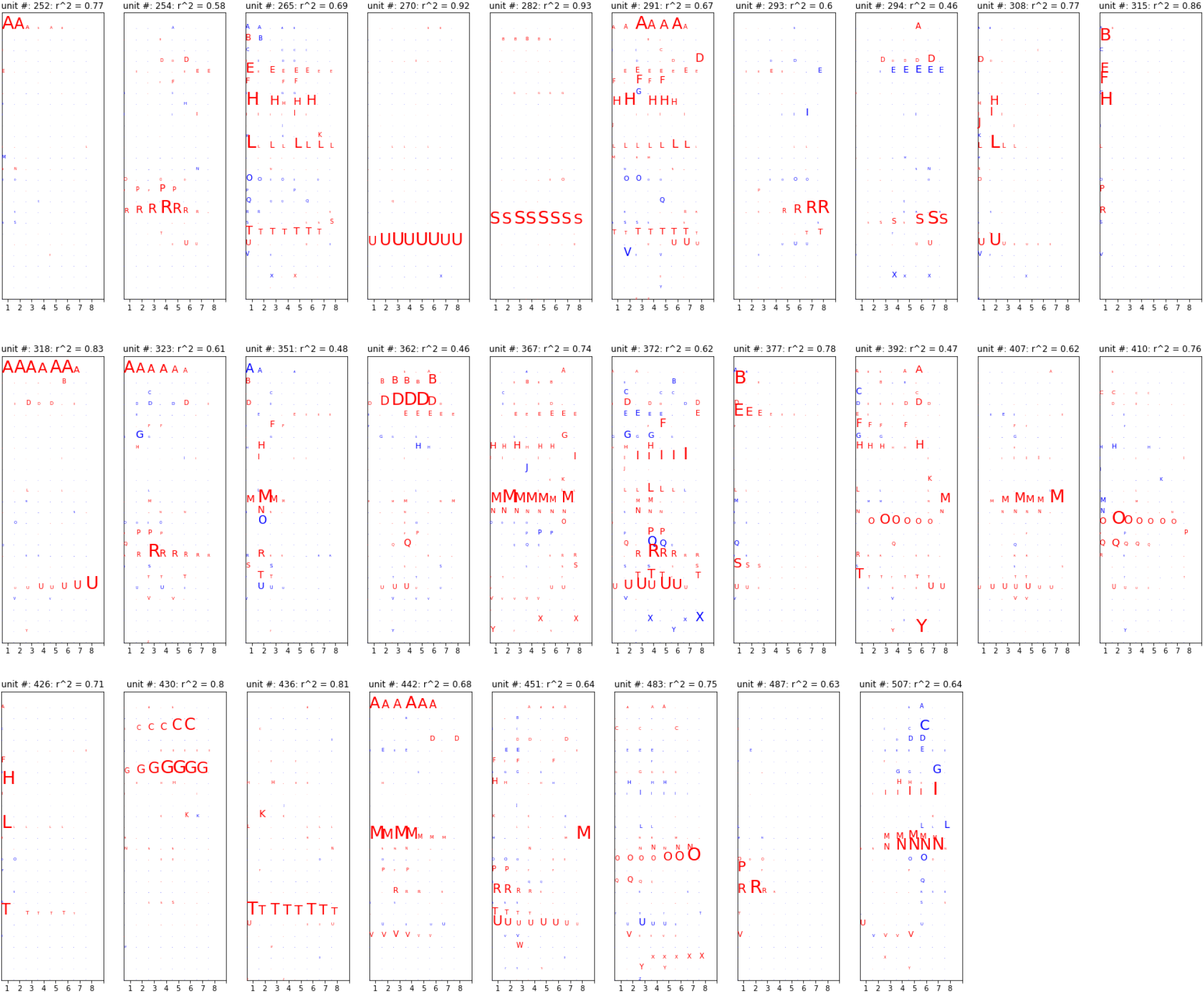
Visualizing the receptive fields of all the word-selective units in the non-biased literate networks

## References

1. Dehaene S, Cohen L, Morais J, Kolinsky R. Illiterate to literate: behavioural and cerebral changes induced by reading acquisition. Nat Rev Neurosci. 2015;16: 234–244. doi:10.1038/nrn3924

2. Wandell BA, Yeatman JD. Biological development of reading circuits. Curr Opin Neurobiol. 2013. doi:10.1016/j.conb.2012.12.005

3. Cohen L, Dehaene S, Naccache L, Lehéricy S, Dehaene-Lambertz G, Hénaff MA, et al. The visual word form area: Spatial and temporal characterization of an initial stage of reading in normal subjects and posterior split-brain patients. Brain. 2000;123: 291–307.

4. Baker CI, Liu J, Wald LL, Kwong KK, Benner T, Kanwisher N. Visual word processing and xperiential origins of functional selectivity in human xtrastriate cortx. Proceedings of the National Academy of Sciences of the United States of America. 2007;104: 9087–92.

5. Szwed M, Qiao E, Jobert A, Dehaene S, Cohen L. Effects of literacy in early visual and occipitotemporal areas of Chinese and French readers. J Cogn Neurosci. 2014;26: 459–475. doi:10.1162/jocn_a_00499

6. Rauschecker AM, Bowen RF, Parvizi J, Wandell BA. Position sensitivity in the visual word form area. Proceedings of the National Academy of Sciences. 2012;109: E1568–E1577. doi:10.1073/pnas.1121304109

7. Dehaene S, Naccache L, Cohen L, Le Bihan D, Mangin JF, Poline JB, et al. Cerebral mechanisms of word masking and unconscious repetition priming. Nat Neurosci. 2001;4: 752–8.

8. Dehaene S, Jobert A, Naccache L, Ciuciu P, Poline JB, Le Bihan D, et al. Letter binding and invariant recognition of masked words: behavioral and neuroimaging evidence. Psychol Sci. 2004;15: 307– 13.

9. Bruno JL, Zumberge A, Manis FR, Lu ZL, Goldman JG. Sensitivity to orthographic familiarity in the occipito-temporal region. Neuroimage. 2008;39: 1988–2001. doi:10.1016/j.neuroimage.2007.10.044

10. Kronbichler M, Hutzler F, Wimmer H, Mair A, Staffen W, Ladurner G. The visual word form area and the frequency with which words are encountered: evidence from a parametric fMRI study. Neuroimage. 2004;21: 946–53. doi:10.1016/j.neuroimage.2003.10.021

11. Dehaene-Lambertz G, Monzalvo K, Dehaene S. The emergence of the visual word form: Longitudinal evolution of category-specific ventral visual areas during reading acquisition. PLoS Biol. 2018;16: e2004103. doi:10.1371/journal.pbio.2004103

12. Price CJ, Devlin JT. The Interactive Account of ventral occipitotemporal contributions to reading. Trends in Cognitive Sciences. 2011;15: 246–253. doi:10.1016/j.tics.2011.04.001

13. Dehaene S, Cohen L. The unique role of the visual word form area in reading. Trends Cogn Sci. 2011;15: 254–62. doi:10.1016/j.tics.2011.04.003

14. Twomey T, Kawabata Duncan KJ, Price CJ, Devlin JT. Top-down modulation of ventral occipito-temporal responses during visual word recognition. NeuroImage. 2011;55: 1242–1251. doi:10.1016/j.neuroimage.2011.01.001

15. Dehaene S, Cohen L. Cultural recycling of cortical maps. Neuron. 2007;56: 384–98.

16. Dehaene S. Reading in the brain. New York: Penguin Viking; 2009.

17. Dehaene S, Pegado F, Braga LW, Ventura P, Nunes Filho G, Jobert A, et al. How learning to read changes the cortical networks for vision and language. Science. 2010;330: 1359–64. doi:10.1126/science.1194140

18. Hasson U, Levy I, Behrmann M, Hendler T, Malach R. Eccentricity bias as an organizing principle for human high-order object areas. Neuron. 2002;34: 479–90.

19. Malach R, Levy I, Hasson U. The topography of high-order human object areas. Trends Cogn Sci. 2002;6: 176–184.

20. Szwed M, Cohen L, Qiao E, Dehaene S. The role of invariant line junctions in object and visual word recognition. Vision Res. 2009;49: 718–25. doi:10.1016/j.visres.2009.01.003

21. Long B, Yu C-P, Konkle T. Mid-level visual features underlie the high-level categorical organization of the ventral stream. PNAS. 2018;115: E9015–E9024. doi:10.1073/pnas.1719616115

22. Barttfeld P, Abboud S, Lagercrantz H, Adén U, Padilla N, Edwards AD, et al. A lateral-to-mesial organization of human ventral visual cortex at birth. Brain Struct Funct. 2018;223: 3107–3119. doi:10.1007/s00429-018-1676-3

23. Bouhali F, Thiebaut de Schotten M, Pinel P, Poupon C, Mangin JF, Dehaene S, et al. Anatomical connections of the visual word form area. J Neurosci. 2014;34: 15402–14. doi:10.1523/JNEUROSCI.4918-13.2014

24. Hannagan T, Amedi A, Cohen L, Dehaene-Lambertz G, Dehaene S. Origins of the specialization for letters and numbers in ventral occipitotemporal cortx. Trends Cogn Sci (Regul Ed). 2015;19: 374–382. doi:10.1016/j.tics.2015.05.006

25. Mahon BZ, Caramazza A. What drives the organization of object knowledge in the brain? Trends Cogn Sci. 2011;15: 97–103. doi:10.1016/j.tics.2011.01.004

26. Mars RB, Sotiropoulos SN, Passingham RE, Sallet J, Verhagen L, Khrapitchev AA, et al. Whole brain comparative anatomy using connectivity blueprints. Stephan KE, editor. eLife. 2018;7: e35237. doi:10.7554/eLife.35237

27. Saygin ZM, Osher DE, Norton ES, Youssoufian DA, Beach SD, Feather J, et al. Connectivity precedes function in the development of the visual word form area. Nature Neuroscience. 2016;19: 1250–1255. doi:10.1038/nn.4354

28. Kubilius J, Schrimpf M, Nayebi A, Bear D, Yamins DLK, DiCarlo JJ. CORnet: Modeling the Neural Mechanisms of Core Object Recognition. bioRxiv. 2018; 408385. doi:10.1101/408385

29. Vinckier F, Dehaene S, Jobert A, Dubus JP, Sigman M, Cohen L. Hierarchical coding of letter strings in the ventral stream: dissecting the inner organization of the visual word-form system. Neuron. 2007;55: 143–56.

30. Binder JR, Medler DA, Westbury CF, Liebenthal E, Buchanan L. Tuning of the human left fusiform gyrus to sublxical orthographic structure. Neuroimage. 2006;33: 739–48.

31. Woolnough O, Donos C, Rollo PS, Forseth KJ, Lakretz Y, Crone NE, et al. Spatiotemporal dynamics of orthographic and lxical processing in the ventral visual pathway. Nature Human Behaviour. 2020; 1–10. doi:10.1038/s41562-020-00982-w

32. Cohen L, Dehaene S, McCormick S, Durant S, Zanker JM. Brain mechanisms of recovery from pure alexia: A single case study with multiple longitudinal scans. Neuropsychologia. 2016;91: 36–49. doi:10.1016/j.neuropsychologia.2016.07.009

33. Dejerine J. Contribution à l’étude anatomo-pathologique et clinique des différentes variétés de cécité verbale. Mem Soc Biol. 1892;4: 61–90.

34. Gaillard R, Naccache L, Pinel P, Clemenceau S, Volle E, Hasboun D, et al. Direct intracranial, FMRI, and lesion evidence for the causal role of left inferotemporal cortex in reading. Neuron. 2006;50: 191–204.

35. Spoerer CJ, Kietzmann TC, Mehrer J, Charest I, Kriegeskorte N. Recurrent neural networks can xplain flxible trading of speed and accuracy in biological vision. PLOS Computational Biology. 2020;16: e1008215. doi:10.1371/journal.pcbi.1008215

36. Glezer LS, Jiang X, Riesenhuber M. Evidence for highly selective neuronal tuning to whole words in the “visual word form area.” Neuron. 2009;62: 199–204. doi:10.1016/j.neuron.2009.03.017

37. Dehaene S, Cohen L, Sigman M, Vinckier F. The neural code for written words: a proposal. Trends Cogn Sci. 2005;9: 335–41.

38. Grainger J, Granier JP, Farioli F, Van Assche E, van Heuven WJ. Letter position information and printed word perception: the relative-position priming constraint. Journal of xperimental psychology. 2006;32: 865–84.

39. Bouhali F, Bézagu Z, Dehaene S, Cohen L. A mesial-to-lateral dissociation for orthographic processing in the visual cortx. PNAS. 2019; 201904184. doi:10.1073/pnas.1904184116

40. Agrawal A, Hari KVS, Arun SP. Reading Increases the Compositionality of Visual Word Representations. Psychol Sci. 2019; 0956797619881134. doi:10.1177/0956797619881134

41. Agrawal A, Hari K, Arun S. A compositional neural code in high-level visual cortexcan xplain jumbled word reading. Baker CI, editor. eLife. 2020;9: e54846. doi:10.7554/eLife.54846

42. Grainger J, Dufau S, Ziegler JC. A Vision of Reading. Trends in Cognitive Sciences. 2016;20: 171– 179. doi:10.1016/j.tics.2015.12.008

43. Rajalingham R, Kar K, Sanghavi S, Dehaene S, DiCarlo JJ. The inferior temporal cortex is a potential cortical precursor of orthographic processing in untrained monkeys. Nature Communications. 2020;11: 3886. doi:10.1038/s41467-020-17714-3

44. Self MW, Peters JC, Possel JK, Reithler J, Goebel R, Ris P, et al. The Effects of Contxt and Attention on Spiking Activity in Human Early Visual Cortx. PLOS Biology. 2016;14: e1002420. doi:10.1371/journal.pbio.1002420

45. Hinton GE, Srivastava N, Krizhevsky A, Sutskever I, Salakhutdinov RR. Improving neural networks by preventing co-adaptation of feature detectors. arXiv:12070580 [cs]. 2012 [cited 21 Jan 2021]. Available: http://arxiv.org/abs/1207.0580

46. Srivastava N, Hinton G, Krizhevsky A, Sutskever I, Salakhutdinov R. Dropout: a simple way to prevent neural networks from overfitting. J Mach Learn Res. 2014;15: 1929–1958.

47. Kriegeskorte N, Mur M, Bandettini PA. Representational similarity analysis-connecting the branches of systems neuroscience. Frontiers in systems neuroscience. 2008;2: 4.

48. Shepard RN. Analysis of proximities as a technique for the study of information processing in man. Hum Factors. 1963;5: 33–48.

49. Shepard RN, Chipman S. Second-order isomorphism of internal representations: Shapes of states. Cognitive Psychology. 1970;1: 1–17.

50. Chang CHC, Pallier C, Wu DH, Nakamura K, Jobert A, Kuo W-J, et al. Adaptation of the human visual system to the statistics of letters and line configurations. Neuroimage. 2015;120: 428–440. doi:10.1016/j.neuroimage.2015.07.028

51. Pegado F, Comerlato E, Ventura F, Jobert A, Nakamura K, Buiatti M, et al. Timing the impact of literacy on visual processing. Proc Natl Acad Sci USA. 2014;111: E5233–5242. doi:10.1073/pnas.1417347111

52. Goodfellow I, Lee H,Le QV, Saxe A, Ng AY. Measuring Invariances in Deep Networks. In: Bengio Y, Schuurmans D, Lafferty JD, CKI Williams, Culotta A, editors. Advances in Neural Information Processing Systems 22. Curran Associates, Inc.; 2009. pp. 646–654. Available: http://papers.nips.cc/paper/3790-measuring-invariances-in-deep-networks.pdf

53. Pflugshaupt T, Gutbrod K, Wurtz P, von Wartburg R, Nyffeler T, de Haan B, et al. About the role of visual field defects in pure alexia. Brain. 2009;132: 1907–17.

54. New B, Ferrand L, Pallier C, Brysbaert M. Rexamineing the word length effect in visual word recognition: new evidence from the English Lxicon Project. Psychon Bull Rev. 2006;13: 45–52.

55. Weekes BS. Differential effects of number of letters on word and nonword naming latency. Quarterly Journal of Xperimental Psychology. 1997;50A: 439–456.

56. Cohen L, Dehaene S, Vinckier F, Jobert A, Montavont A. Reading normal and degraded words: contribution of the dorsal and ventral visual pathways. NeuroImage. 2008;40: 353–66.

57. Aghababian V, Nazir TA. Developing normal reading skills: aspects of the visual processes underlying word recognition. J Xp Child Psychol. 2000;76: 123–50.

58. Cohen L, Martinaud O, Lemer C, Lehéricy S, Samson Y, Obadia M, et al. Visual word recognition in the left and right hemispheres: Anatomical and functional correlates of peripheral alexias. Cerebral Cortx. 2003;13: 1313–1333.

59. Graves WW, Desai R, Humphries C, Seidenberg MS, Binder JR. Neural systems for reading aloud: a multiparametric approach. Cereb Cortx. 2010;20: 1799–815. doi:10.1093/cercor/bhp245

60. Coltheart M, Rastle K, Perry C, Langdon R, Ziegler J. DRC: a dual route cascaded model of visual word recognition and reading aloud. Psychol Rev. 2001;108: 204–56.

61. Harm MW, Seidenberg MS. Phonology, reading acquisition, and dyslxia: insights from connectionist models. Psychol Rev. 1999;106: 491–528.

62. Seidenberg MS, McClelland JL. A distributed, developmental model of word recognition and naming. Psychol Rev. 1989;96: 523–68.

63. Zorzi M, Houghton G, Butterworth B. Two routes or one in reading aloud? A connectionist dual-process model. Journal of Experimental Psychology: Human Perception and Performance. 1998;24: 1131–1161.

64. Plaut DC, McClelland JL, Seidenberg MS, Patterson K. Understanding normal and impaired word reading: computational principles in quasi-regular domains. Psychol Rev. 1996;103: 56–115.

65. McCoy RT, Linzen T, Dunbar E, Smolensky P. RNNs Implicitly Implement Tensor Product Representations. arXiv:181208718 [cs]. 2019 [cited 28 Mar 2020]. Available: http://arxiv.org/abs/1812.08718

66. Smolensky P. Tensor Product Variable Binding and the Representation of Symbolic Structures in Connectionist Systems. Artificial Intelligence. 1990;46: 159–216.

67. McCloskey M, Cohen NJ. Catastrophic Interference in Connectionist Networks: The Sequential Learning Problem. In: ower GH, editor. Psychology of Learning and Motivation. Academic Press; 1989. pp. 109–165. doi:10.1016/S0079-7421(08)60536-8

68. Kolinsky R. How Learning to Read Influences Language and Cognition. In: Pollatsek A, Treiman R, editors. The Oxford Handbook of Reading. Oxford University Press; 2014. p. Available on-line at www.oxfordhandbooks.com.

69. Kubota EC, Joo SJ, Huber E, Yeatman JD. Word selectivity in high-level visual cortex and reading skill. Developmental Cognitive Neuroscience. 2019;36: 100593. doi:10.1016/j.dcn.2018.09.003

70. Pegado F, Nakamura K, Cohen L, Dehaene S. Breaking the symmetry: mirror discrimination for single letters but not for pictures in the Visual Word Form Area. Neuroimage. 2011;55: 742–9. doi:10.1016/j.neuroimage.2010.11.043

71. Pegado F, Nakamura K, Braga LW, Ventura P, Nunes Filho G, Pallier C, et al. Literacy breaks mirror invariance for visual stimuli: a behavioral study with adult illiterates. J Xp Psychol Gen. 2014;143: 887–894. doi:10.1037/a0033198

72. Chang L, Tsao DY. The Code for Facial Identity in the Primate Brain. Cell. 2017;169: 1013-1028.e14. doi:10.1016/j.cell.2017.05.011

73. Miller GA, Chomsky N. Finitary Models of Language Users1. Handbook of Mathematical Psychology: VOLUME 1-CHAPTERS 1-8. 1963; 419.

74. Vinckier F, Qiao E, Pallier C, Dehaene S, Cohen L. The impact of letter spacing on reading: A test of the bigram coding hypothesis. J Vis. 2011;11. doi:10.1167/11.6.8

75. Lee H, Margalit E, Jozwik KM, Cohen MA, Kanwisher N, Yamins DL, et al. Topographic deep artificial neural networks reproduce the hallmarks of the primate inferior temporal cortex face processing network. bioRxiv. 2020.

76. Schrimpf M, Kubilius J, Hong H, Majaj NJ, Rajalingham R, Issa EB, et al. Brain-Score: Which Artificial Neural Network for Object Recognition is most Brain-Like? bioRxiv. 2018; 407007. doi:10.1101/407007

77. Kietzmann TC, Spoerer CJ, Sörensen LKA, Cichy RM, Hauk O, Kriegeskorte N. Recurrence is required to capture the representational dynamics of the human visual system. PNAS. 2019;116: 21854–21863. doi:10.1073/pnas.1905544116

78. Yamins DL, Hong H, Cadieu CF, Solomon EA, Seibert D, DiCarlo JJ. Performance-optimized hierarchical models predict neural responses in higher visual cortx. Proceedings of the National Academy of Sciences. 2014;111: 8619–8624.

79. Eickenberg M, Gramfort A, Varoquaux G, Thirion B. Seeing it all: Convolutional network layers map the function of the human visual system. Neuroimage. 2017;152: 184–194. doi:10.1016/j.neuroimage.2016.10.001

80. Bao P, She L, McGill M, Tsao DY. A map of object space in primate inferotemporal cortx. Nature. 2020; 1–6. doi:10.1038/s41586-020-2350-5

81. Nayebi A, Bear D, Kubilius J, Kar K, Ganguli S, Sussillo D, et al. Task-Driven Convolutional Recurrent Models of the Visual System. arXiv:180700053 [cs, q-bio]. 2018 [cited 10 Feb 2021]. Available: http://arxiv.org/abs/1807.00053

82. Melby-Lervåg M, Lyster S-AH, Hulme C. Phonological skills and their role in learning to read: a meta-analytic review. Psychol Bull. 2012;138: 322–352. doi:10.1037/a0026744

83. Castles A, Rastle K, Nation K. Ending the Reading Wars: Reading Acquisition From Novice to Xpert. Psychol Sci Public Interest. 2018;19: 5–51. doi:10.1177/1529100618772271

84. Di Bono MG, Zorzi M. Deep generative learning of location-invariant visual word recognition. Front Psychol. 2013;4: 635. doi:10.3389/fpsyg.2013.00635

85. Jaderberg M, Simonyan K, Vedaldi A, Zisserman A. Reading Txt in the Wild with Convolutional Neural Networks. Int J Comput Vis. 2016;116: 1–20. doi:10.1007/s11263-015-0823-z

86. Grainger J, Dufau S, Montant M, Ziegler JC, Fagot J. Orthographic Processing in Baboons (Papio papio). Science. 2012;336: 245–248. doi:10.1126/science.1218152

87. Srihasam K, Mandeville JB, Morocz IA, Sullivan KJ, Livingstone MS. Behavioral and Anatomical Consequences of Early versus Late Symbol Training in Macaques. Neuron. 2012;73: 608–619. doi:10.1016/j.neuron.2011.12.022

88. Tsao DY, Freiwald WA, Tootell RB, Livingstone MS. A cortical region consisting entirely of face-selective cells. Science. 2006;311: 670–4.

